## Supplementary Figures for "Learning from the expert: studying *Salicornia* to understand salinity tolerance"

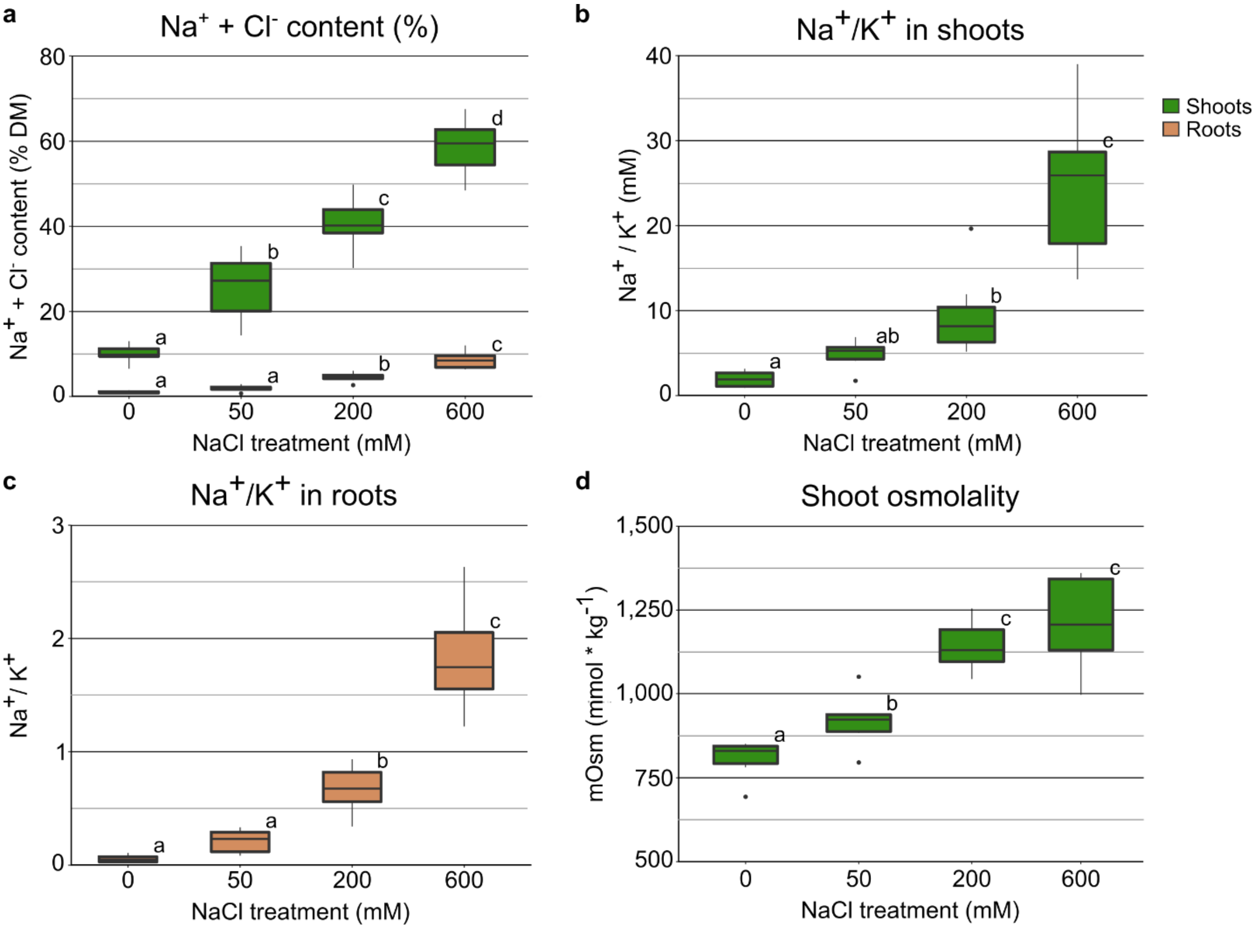


**Supplementary Figure 1.** Ion and osmolyte accumulation in *S. bigelovii* treated with 0, 50, 200, or 600 mM NaCl. **a,** Sodium and chloride content as percentage of dry mass. **b,** Sodium and potassium ratios in shoots. **c,** Sodium and potassium ratios in roots. **d,** Shoot osmolality. Mean differences were compared and the FDR was controlled with the Benjamini-Hochberg procedure at an α = 0.05, significant differences are indicated as different letters.

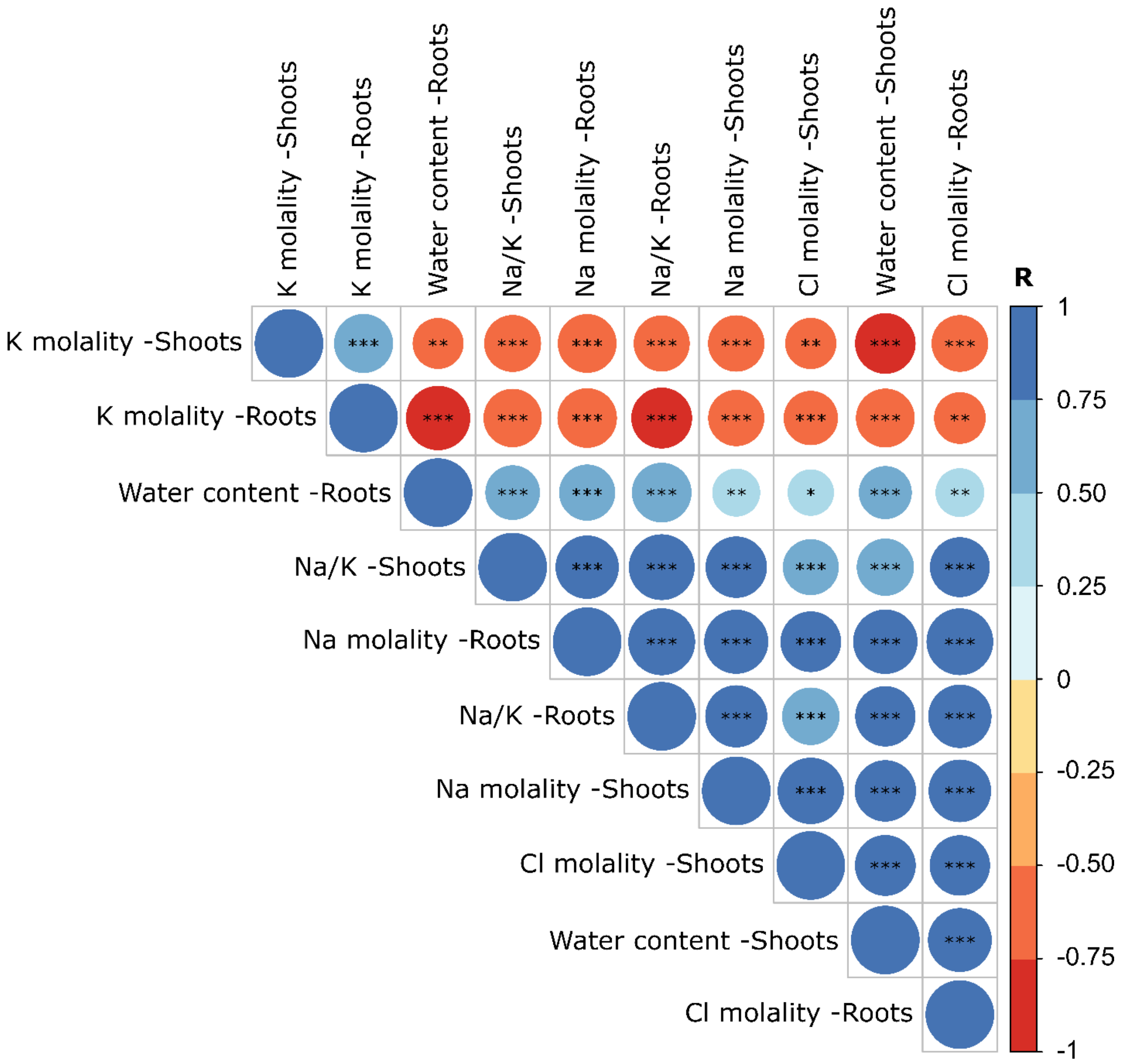


**Supplementary Figure 2.** Correlations between ion accumulation and water content in *S. bigelovii* plants treated with 0, 50, 200, or 600 mM NaCl. Correlation coefficients are displayed as color and size gradients between positive (blue) and negative (red) values. To control the FDR, P-values were corrected with the Benjamini-Hochberg procedure and depicted with different levels of significance as: *, 0.05; **, 0.01; and ***, 0.001.

#### *S. bigelovii*

**a** **b**

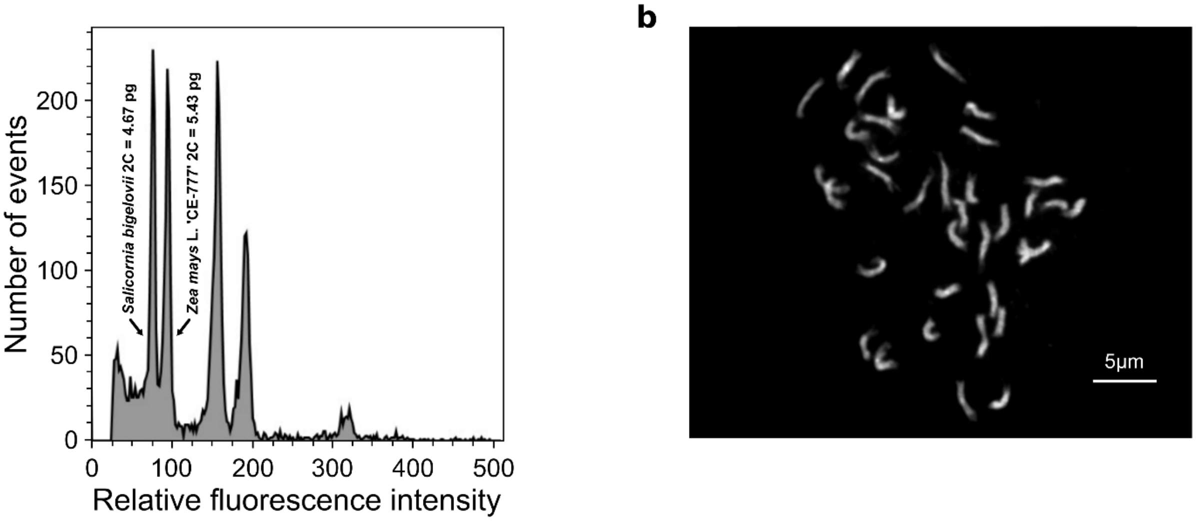


**c**

**Genome size estimations**

| Method | Genome size estimation |
|---|---|
| Flow cytometry | 2,138,000,000 bp |
| FindGSE (Illumina) | 2,015,124,775 bp |
| FindGSE (HiFi) | 1,945,431,122 bp |

**d**

**Assembly completeness with Merqury**

| Source | Number of *k*-mers |
|---|---|
| Genome | 650,804,451 |
| Reads | 670,014,626 |
| Percentage of read *k*-mers in the genome | 97.13% |

**Supplementary Figure 3.** *S. bigelovii* genome size estimations and assembly completeness. **a,** Nuclear genome size estimation based on flow cytometry through propidium iodide nuclei staining of *S. bigelovii* and *Zea mays* L 'CE-777' as reference. **b,** Mitotic phase plate of *S. bigelovii* showing 36 chromosomes (2n = 4x = 36). Chromosomes were counterstained with DAPI (light grey pseudocolor). **c,** *S. bigelovii* genome size estimations with flow cytometry and FindGSE. **d,** Assembly completeness with Merqury by comparing the different *k*-mers in the reads and the assembled genome.

### *S. europaea*

**a** **b**

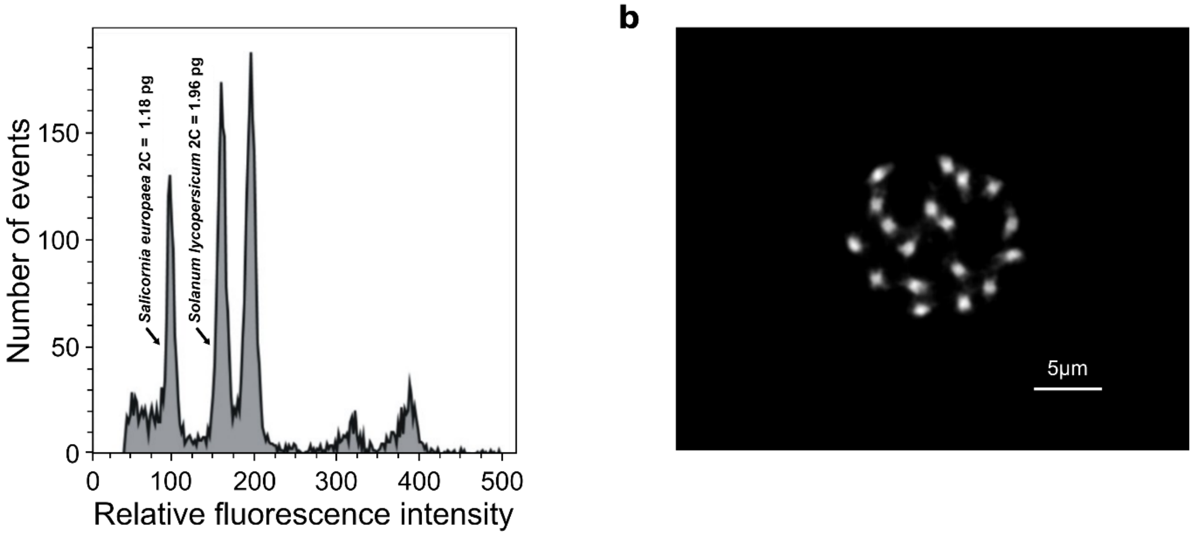


**c**

**Genome size estimations**

| Method | Genome size estimation |
|---|---|
| Flow cytometry | 580,000,000 bp |
| FindGSE (Illumina) | 573,063,611 bp |
| FindGSE (HiFi) | 510,685,893 bp |

**d**

**Assembly completeness with Merqury**

| Source | Number of *k*-mers |
|---|---|
| Genome | 309,298,290 |
| Reads | 311,507,157 |
| Percentage of read *k*-mers in the genome | 99.29% |

**Supplementary Figure 4.** *S. europaea* genome size estimations and assembly completeness. **a,** Nuclear genome size estimation based on flow cytometry through propidium iodide nuclei staining of *S. europaea* and *Solanum lycopersicum* L. ‘Stupické polní rané’ as reference. **b,** Mitotic phase plate of *S. europaea* showing 18 chromosomes (2n = 2x = 18). Chromosomes were counterstained with DAPI (light grey pseudocolor). **c,** *S. europaea* genome size estimations with flow cytometry and FindGSE. **d,** Assembly completeness with Merqury by comparing the different *k*-mers in the reads and the assembled genome.

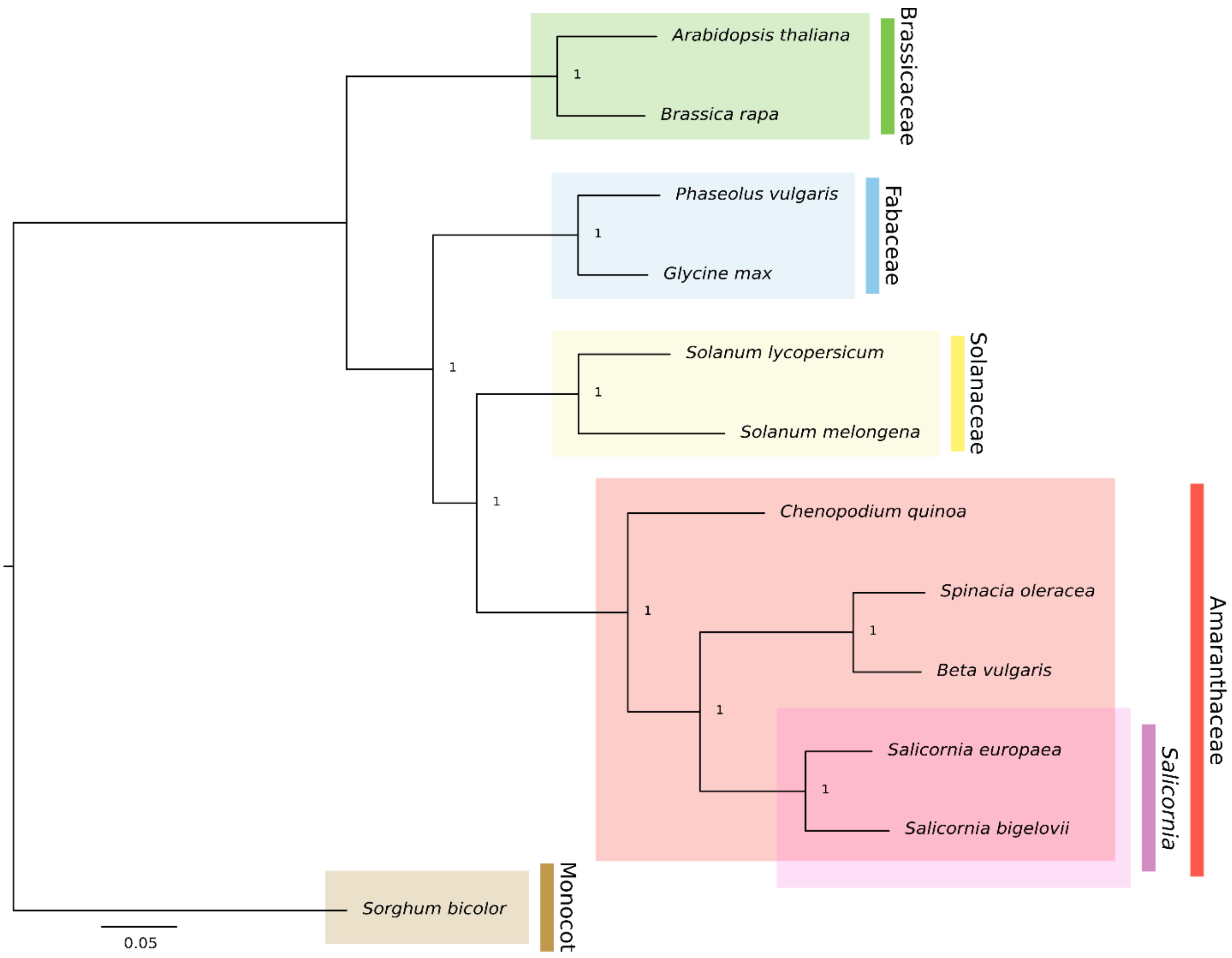


**Supplementary Figure 5.** Phylogenetic reconstruction of the plant families used in this study. Maximum likelihood phylogenetic reconstruction based on 1,377 orthologous proteins selected from the BUSCO Eudicotyledons dataset and found in every species. Values at branching nodes represent transfer bootstrap expectation (TBE) values based on 1,000 replicates. Green, Brassicaceae; blue, Fabaceae; yellow, Solanaceae; red, Amaranthaceae; purple, *Salicornia*; brown, Poaceae (a monocotyledon outgroup).

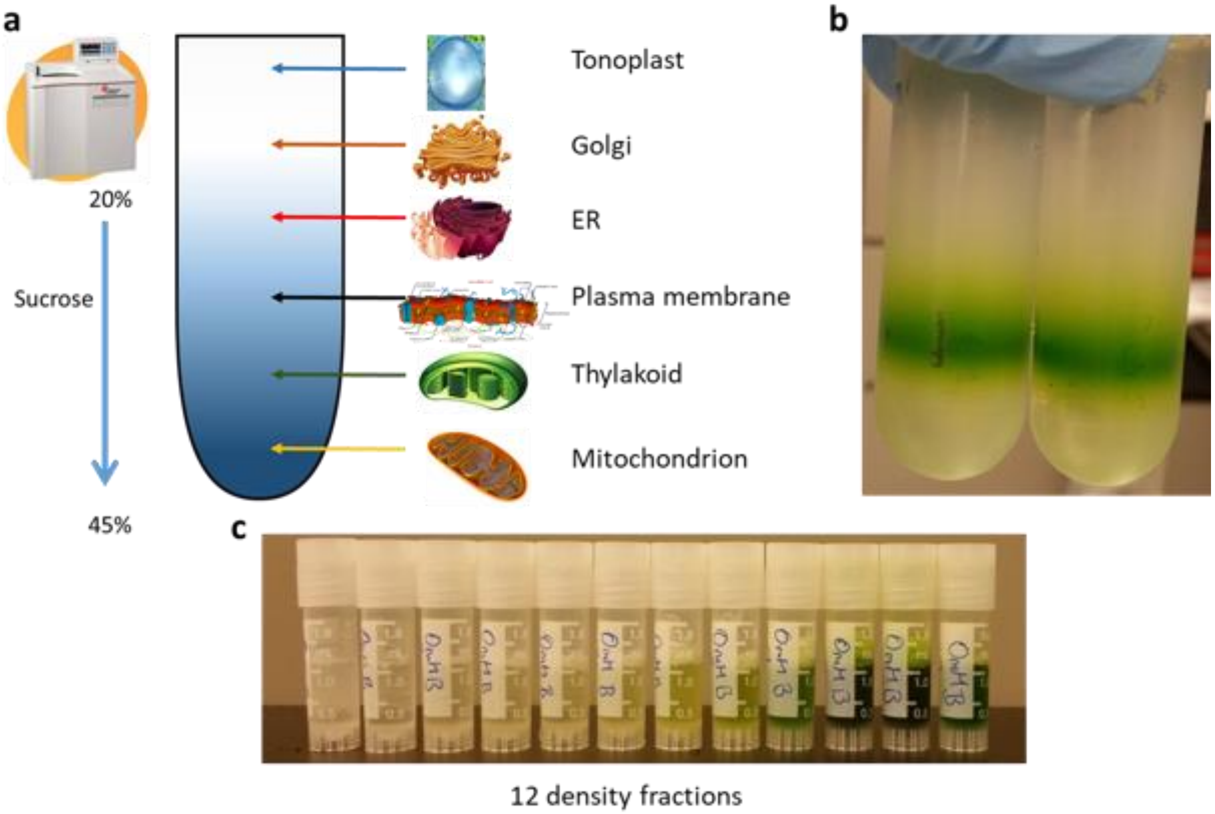


**Supplementary Figure 6.** Separation of organellar membranes by their densities through ultracentrifugation. **a,** Representation of the classical separation of organellar membranes by a continuous sucrose density gradient from 20 - 45% through ultracentrifugation. **b,** Continuous sucrose density gradient of *S. bigelovii* shoot microsomal fraction after ultracentrifugation. **c,** The recovered 12 fractions from *S. bigelovii* shoot microsomal fraction from the continuous sucrose density gradient, each enriched with membranes from different organelles.

**a**

| | Treatment | | | | | | | | | | | | |
|---|---|---|---|---|---|---|---|---|---|---|---|---|---|
| Fraction | 0-A | 0-B | 0-C | 50-A | 50-B | 50-C | 200-A | 200-B | 200-C | 600-A | 600-B | 600-C | Specific gravity range |
| 1 | 19.7 | 19.7 | 20.2 | 19.7 | 19.9 | 20.1 | 20.2 | 20.1 | 20.2 | 20.1 | 20.3 | 19.9 | 1. 08164-1.08431 |
| 2 | 22.4 | 22.5 | 22.4 | 22.4 | 22.4 | 21.9 | 22.3 | 22.5 | 22.7 | 22.5 | 22.5 | 22.2 | 1.09149-1 .09511 |
| 3 | 24.1 | 24 | 23.9 | 23.9 | 23.7 | 23.6 | 23.9 | 24.1 | 24.3 | 24 | 24.1 | 23.9 | 1.09921-1.10241 |
| 4 | 25.5 | 25.5 | 25.4 | 25.3 | 25.2 | 25.4 | 25.5 | 25.8 | 25.8 | 25.6 | 25.7 | 25.4 | 1.10656-1.10934 |
| 5 | 27.3 | 27.5 | 27.4 | 27 | 27.1 | 27.2 | 27.3 | 27.8 | 27.7 | 27.4 | 27.4 | 27.2 | 1.11493-1.11869 |
| 6 | 29.5 | 29.6 | 29.5 | 29.2 | 29.2 | 29.3 | 29.4 | 30 | 29.8 | 29.6 | 29.5 | 29.3 | 1.12532-1.12913 |
| 7 | 31.5 | 31.7 | 31.6 | 31.4 | 31.4 | 31.5 | 31.5 | 32.1 | 31.9 | 31.6 | 31.6 | 31.5 | 1 .13587-1.13926 |
| 8 | 33.7 | 33.9 | 33.8 | 33.5 | 33.6 | 33.7 | 33.7 | 34.2 | 34.1 | 33.8 | 33.7 | 33.6 | 1.14609-1.14954 |
| 9 | 35.8 | 36.2 | 36.1 | 35.7 | 35.9 | 35.6 | 35.9 | 36.3 | 36.3 | 36 | 35.9 | 35.9 | 1.15648-1.15997 |
| 10 | 38 | 38.3 | 38.2 | 37.9 | 38.1 | 37.9 | 38.1 | 38.4 | 38.3 | 38.1 | 38.1 | 38 | 1.16803-1.17056 |
| 11 | 40.1 | 40.5 | 40.3 | 40 | 40.2 | 40.1 | 40.2 | 40.4 | 40.4 | 40.3 | 40.3 | 40.1 | 1.17874-1.18132 |
| 12 | 42.4 | 42.8 | 42.5 | 42.5 | 42.4 | 42.4 | 42.8 | 42.9 | 42.8 | 42.6 | 42.5 | 42.4 | 1.19119-1.19381 |

**b**

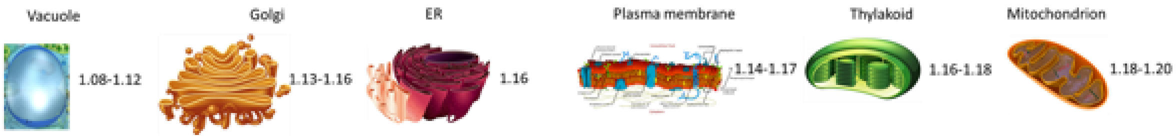


**Supplementary Figure 7.** Composition of the 12 density fractions. **a,** Sucrose % and specific gravity of each of the 12 density fractions recovered from shoots of *S. bigelovii* plants treated with 0, 50, 200, and 600 mM NaCl for 6 weeks. **b,** Specific gravity of different organellar membranes from *Arabidopsis thaliana*. Specific gravity is defined as the substance density relative to water.

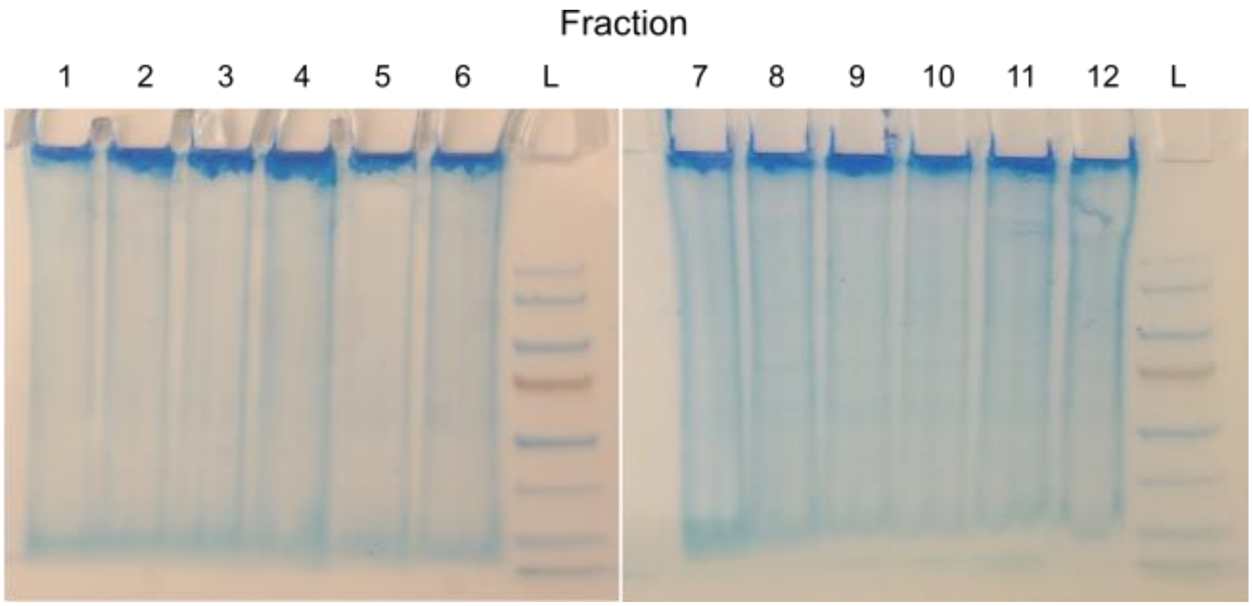


**Supplementary Figure 8.** SDS PAGE of the 12 membrane enriched density fractions of *S. bigelovii* to ensure equal protein loading for subsequent western blot. Fractions (1-12) are numbered according to their relative densities. Ladder is shown as L. 50 µg of protein were loaded per fraction and the gel was stained with Coomassie blue.

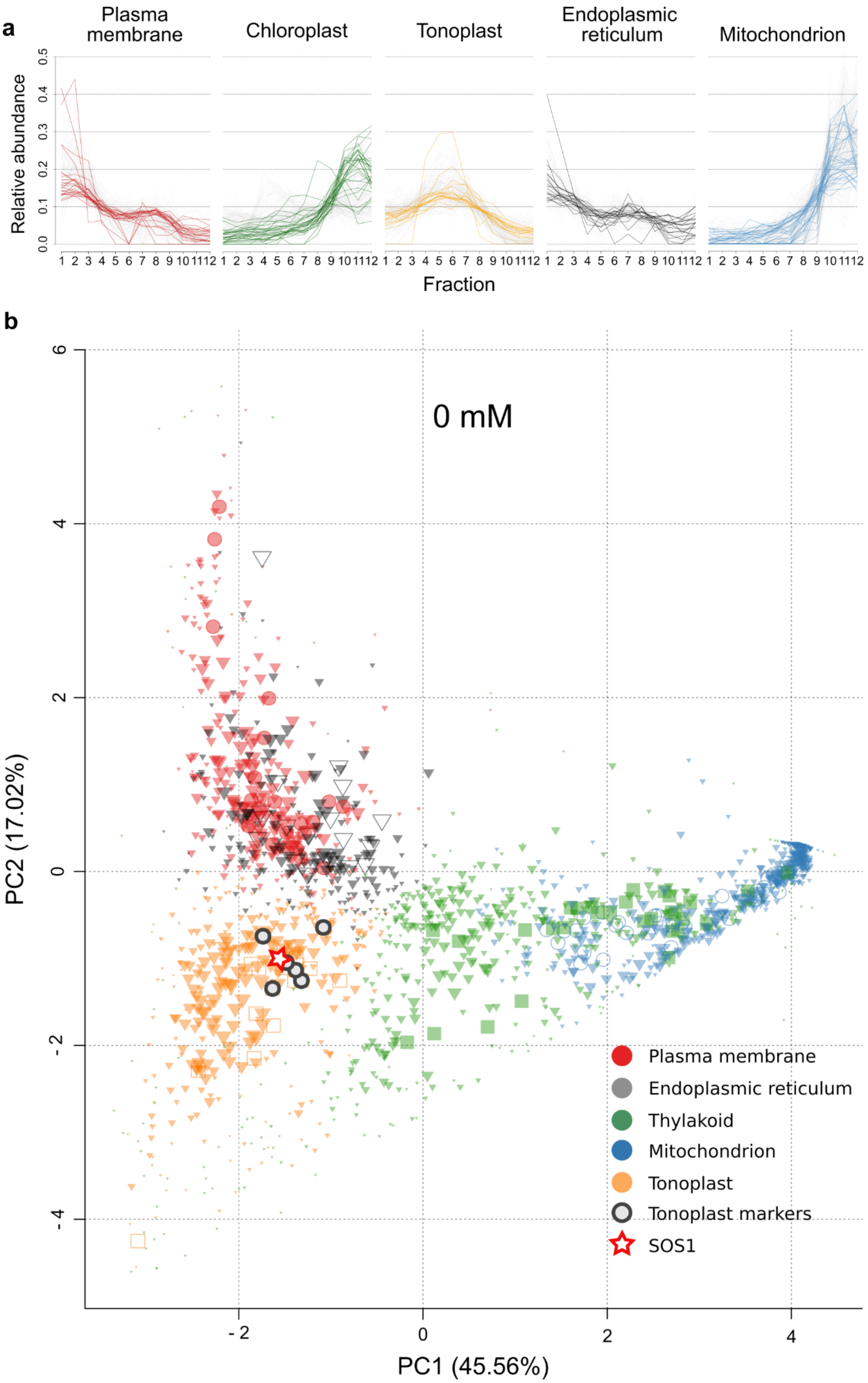
a
Plasma membrane
Chloroplast
Tonoplast
Endoplasmic reticulum
Mitochondrion
Relative abundance
Fraction
b
0 mM
PC2 (17.02%)
PC1 (45.56%)
Plasma membrane
Endoplasmic reticulum
Thylakoid
Mitochondrion
Tonoplast
Tonoplast markers
SOS1

**Supplementary Figure 9.** Spatial protein profiles generated with pRoloc from membrane preparations from shoots of *S. bigelovii* plants treated with 0 mM NaCl. **a,** Relative protein abundance profiles identified with pRoloc of organelle marker proteins on the 12 density fractions. **b,** Principal component analysis of protein predicted subcellular localization by pRoloc. The size of the spot represents confidence values of localization. Plasma membrane: red, markers as circles; endoplasmic reticulum: grey, markers as open triangles; thylakoid: green, markers as squares; mitochondrion: blue, markers as open circles; tonoplast: orange, markers as open squares; tonoplast marker proteins, encircled in black: Vacuolar ATPase subunits a, b, c and d, Vacuolar pyrophosphatase, and Tonoplast Intrinsic Protein 1-3; and red star, SOS1.

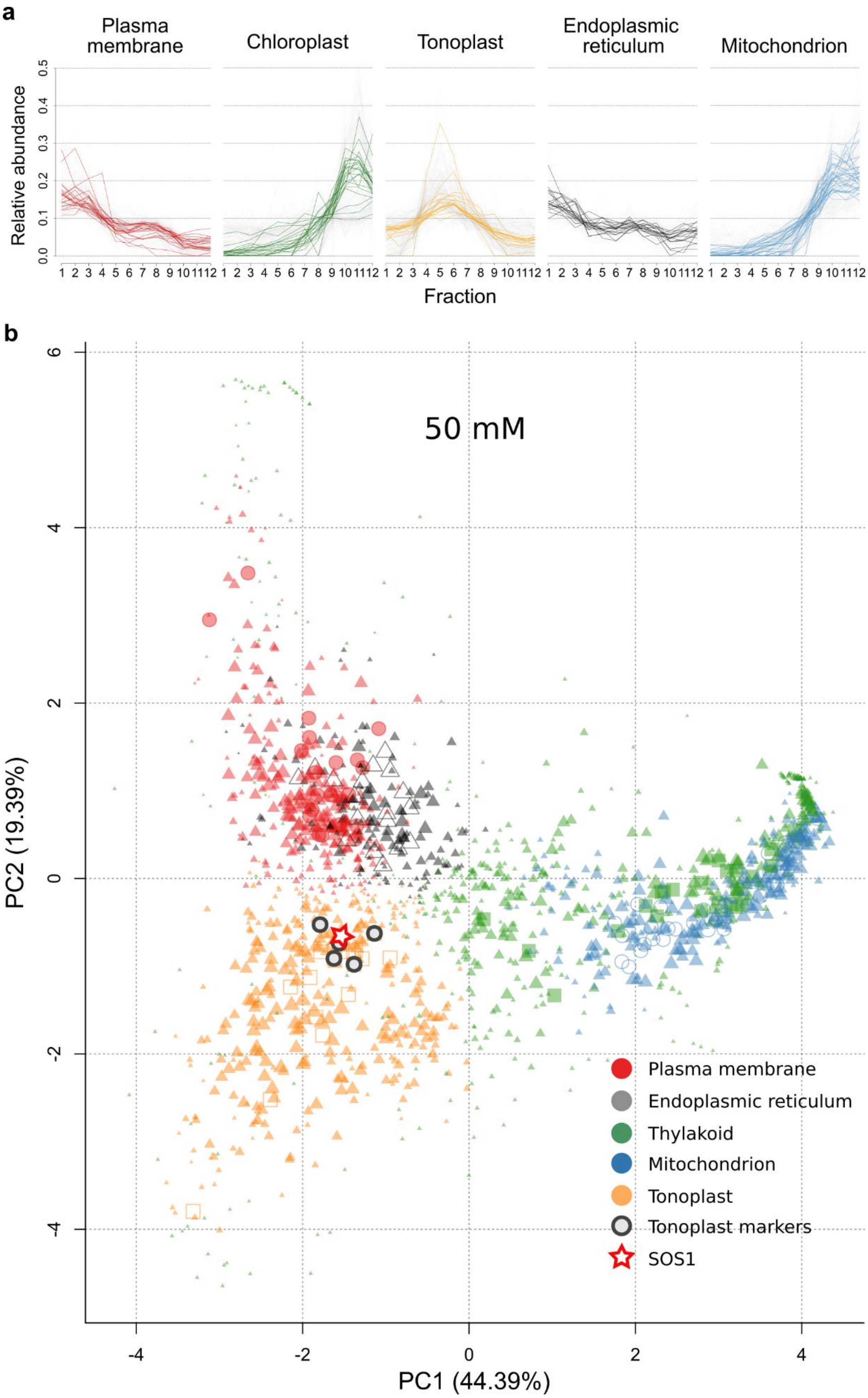
a
Plasma membrane
Chloroplast
Tonoplast
Endoplasmic reticulum
Mitochondrion
Relative abundance
0.0
0.1
0.2
0.3
0.4
0.5
1 2 3 4 5 6 7 8 9 10 11 12
Fraction
b
50 mM
PC2 (19.39%)
PC1 (44.39%)
-4
-2
0
2
4
6
Plasma membrane
Endoplasmic reticulum
Thylakoid
Mitochondrion
Tonoplast
Tonoplast markers
SOS1

**Supplementary Figure 10.** Spatial protein profiles generated with pRoloc from membrane preparations from shoots of *S. bigelovii* plants treated with 50 mM NaCl. **a,** Relative protein abundance profiles identified with pRoloc of organelle marker proteins on the 12 density fractions. **b,** Principal component analysis of protein predicted subcellular localization by pRoloc. The size of the spot represents confidence values of localization. Plasma membrane: red, markers as circles; endoplasmic reticulum: grey, markers as open triangles; thylakoid: green, markers as squares; mitochondrion: blue, markers as open circles; tonoplast: orange, markers as open squares; tonoplast marker proteins, encircled in black: Vacuolar ATPase subunits a, b, c and d, Vacuolar pyrophosphatase, and Tonoplast Intrinsic Protein 1-3; and red star, SOS1.

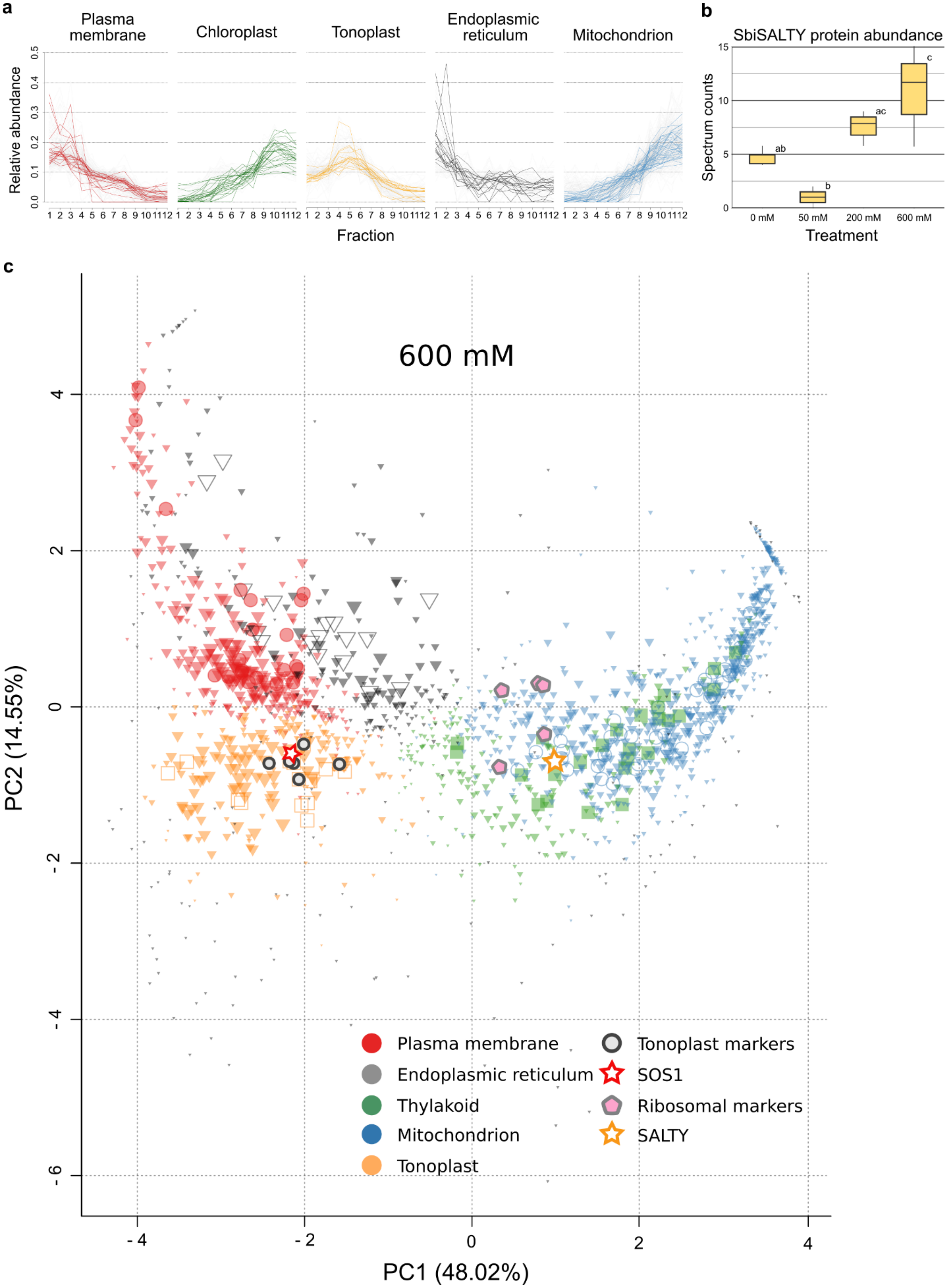

a
Plasma membrane
Chloroplast
Tonoplast
Endoplasmic reticulum
Mitochondrion
Relative abundance
0.0
0.1
0.2
0.3
0.4
0.5
1 2 3 4 5 6 7 8 9 10 11 12
Fraction
b
SbiSALTY protein abundance
Spectrum counts
0
5
10
15
ab
b
ac
c
0 mM
50 mM
200 mM
600 mM
Treatment
c
600 mM
PC2 (14.55%)
PC1 (48.02%)
4
2
0
-2
-4
-6
Plasma membrane
Endoplasmic reticulum
Thylakoid
Mitochondrion
Tonoplast
Tonoplast markers
SOS1
Ribosomal markers
SALTY

**Supplementary Figure 11.** Spatial protein profiles generated with pRoloc from membrane preparations from shoots of *S. bigelovii* plants treated with 600 mM NaCl. **a,** Relative protein abundance profiles identified with pRoloc of organelle marker proteins on the 12 density fractions. **b,** SbiSALTY protein abundance based on spectrum counts per treatment. Mean differences were compared and the FDR was controlled with the Benjamini-Hochberg procedure at an $\alpha = 0.05$, significant differences are indicated as different letters. **c,** Principal component analysis of protein predicted subcellular localization by pRoloc. The size of the spot represents confidence values of localization. Plasma membrane: red, markers as circles; endoplasmic reticulum: grey, markers as open triangles; thylakoid: green, markers as squares; mitochondrion: blue, markers as open circles; tonoplast: orange, markers as open squares; tonoplast marker proteins, encircled in black: Vacuolar ATPase subunits a, b, c and d, Vacuolar pyrophosphatase, and Tonoplast Intrinsic Protein 1-3; red star, SOS1; lilac pentagons: 40S ribosomal protein S7 and 60S ribosomal proteins L14-1, L18a, and L35 ;and yellow star, SALTY.

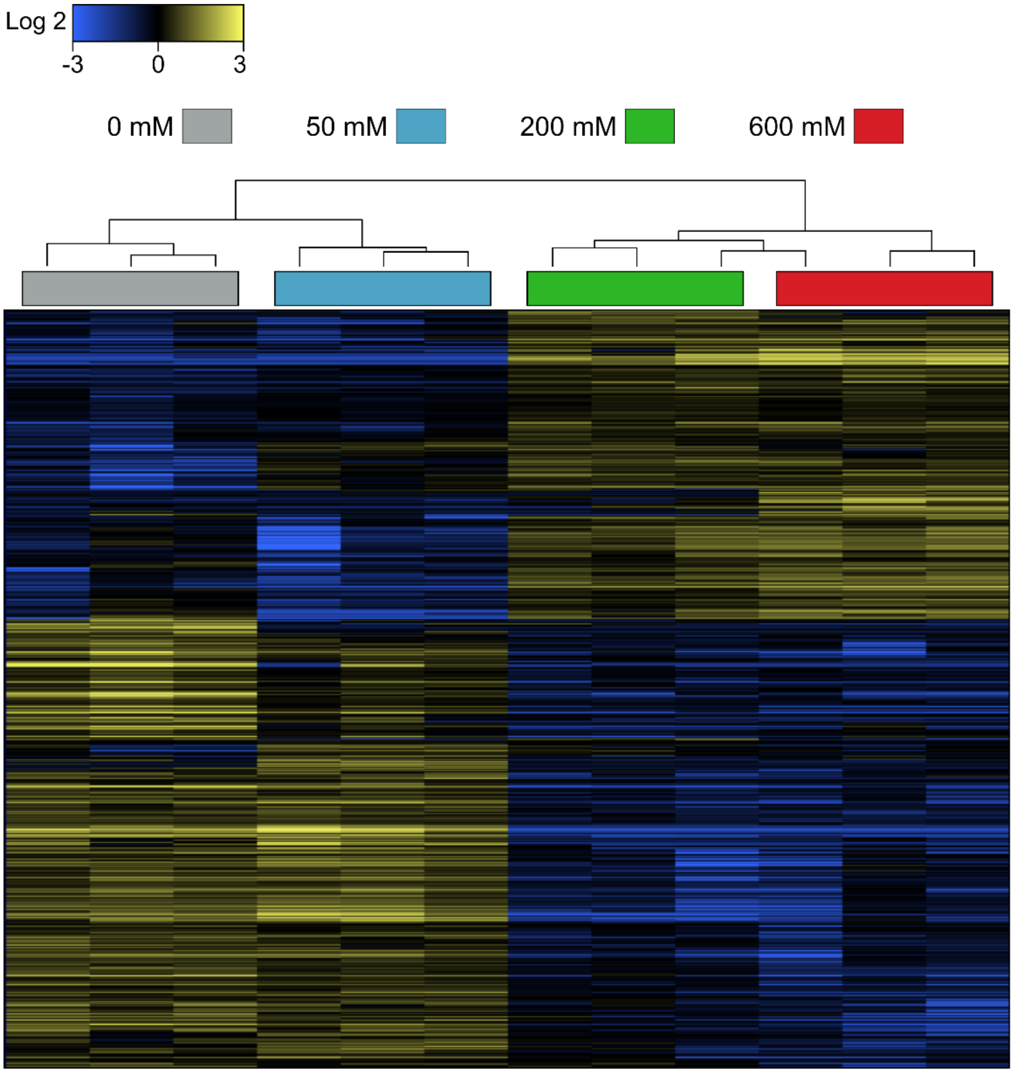


| 0 mM | 50 mM | 200 mM | 600 mM | Number of proteins |
|---|---|---|---|---|
| high | high | low | low | 286 |
| low | low | high | high | 196 |
| high | low | low | low | 71 |
| low | high | low | low | 35 |
| low | high | high | high | 32 |
| low | low | low | high | 31 |
| low | low | high | low | 8 |
| low | high | high | low | 7 |
| high | low | high | high | 7 |
| high | high | high | low | 6 |
| high | low | low | high | 3 |
| high | low | high | low | 2 |
| high | high | low | high | 2 |
| low | high | low | high | 1 |
| | | | | Total: 687 |

**Supplementary Figure 12.** Differential abundance of proteins in *S. bigelovii* shoots. **a,** Hierarchical clustering of three replicates per treatment of differentially abundant proteins in shoots *of S. bigelovii* plants treated with 0, 50, 200, and 600 mM NaCl for 6 weeks. 687 protein clusters had significantly different abundances in at least one treatment. Blue, reduced abundance; Yellow, increased abundance. **b,** Classification of proteins based on their differential abundances and their directionality across treatments, identified with SCAFFOLD.

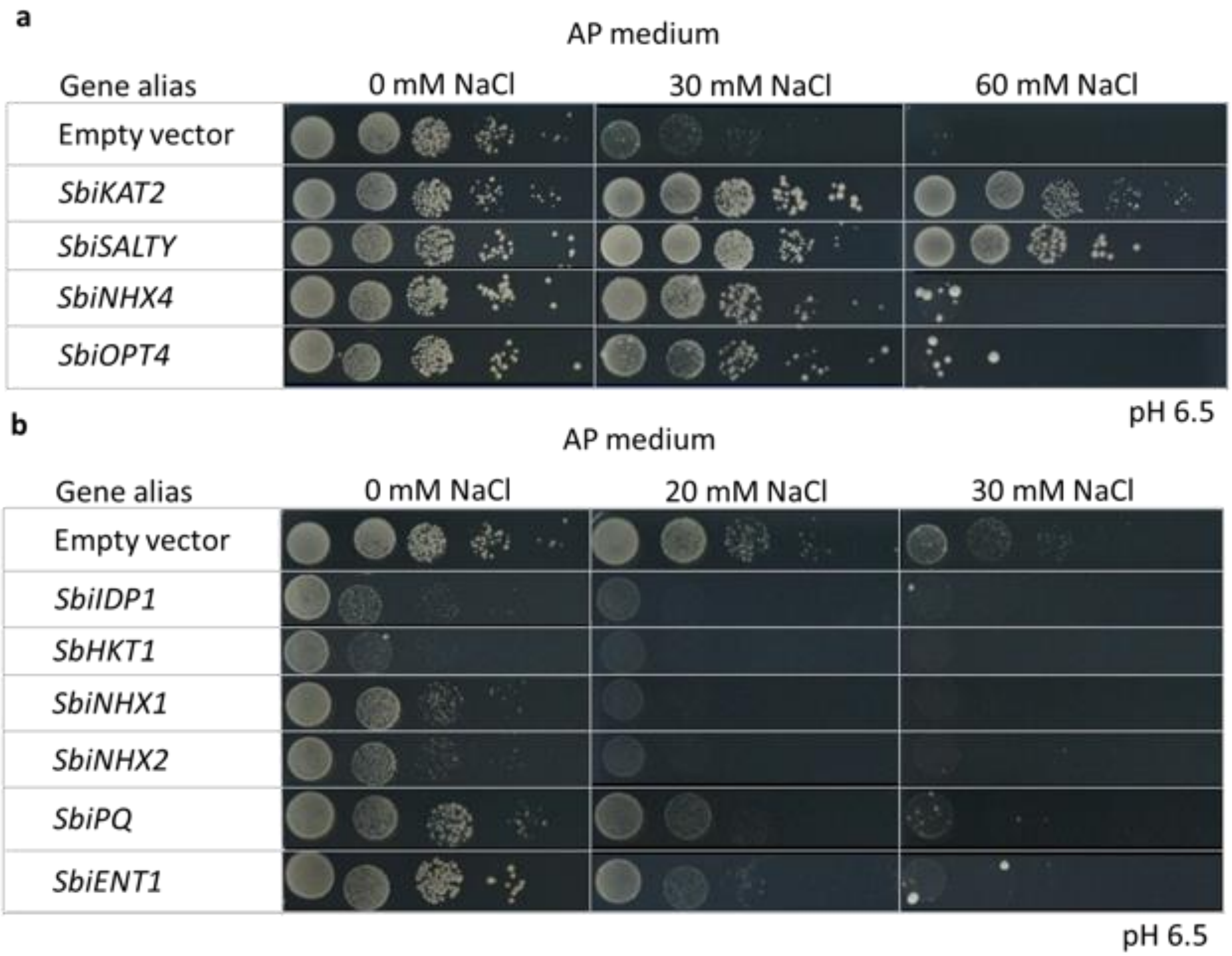


**Supplementary Figure 13.** Yeast spot assays in AXT3 strain at different salt concentrations. **a,** genes increasing salt tolerance. **b,** genes decreasing salt tolerance. Assays were done in AP medium pH 6.5 and supplemented with 2% galactose for gene induction.

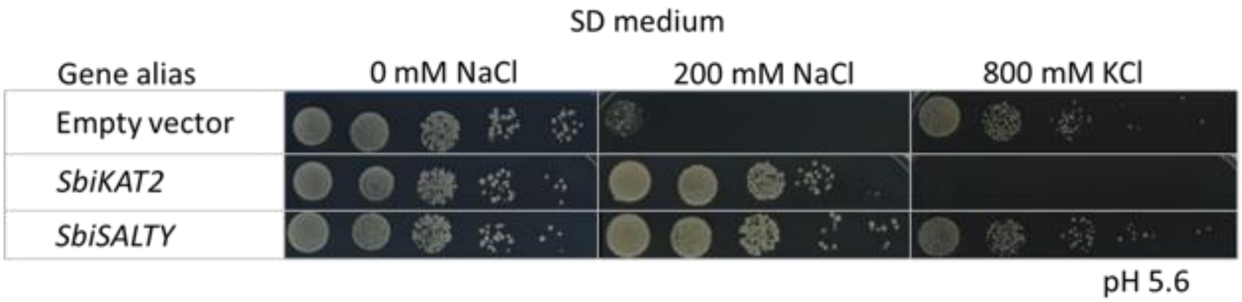


**Supplementary Figure 14.** Yeast spot assays of in AXT3 under high NaCl and KCl. Assays were done in SD medium pH 5.6 and supplemented with 2% galactose for gene induction.

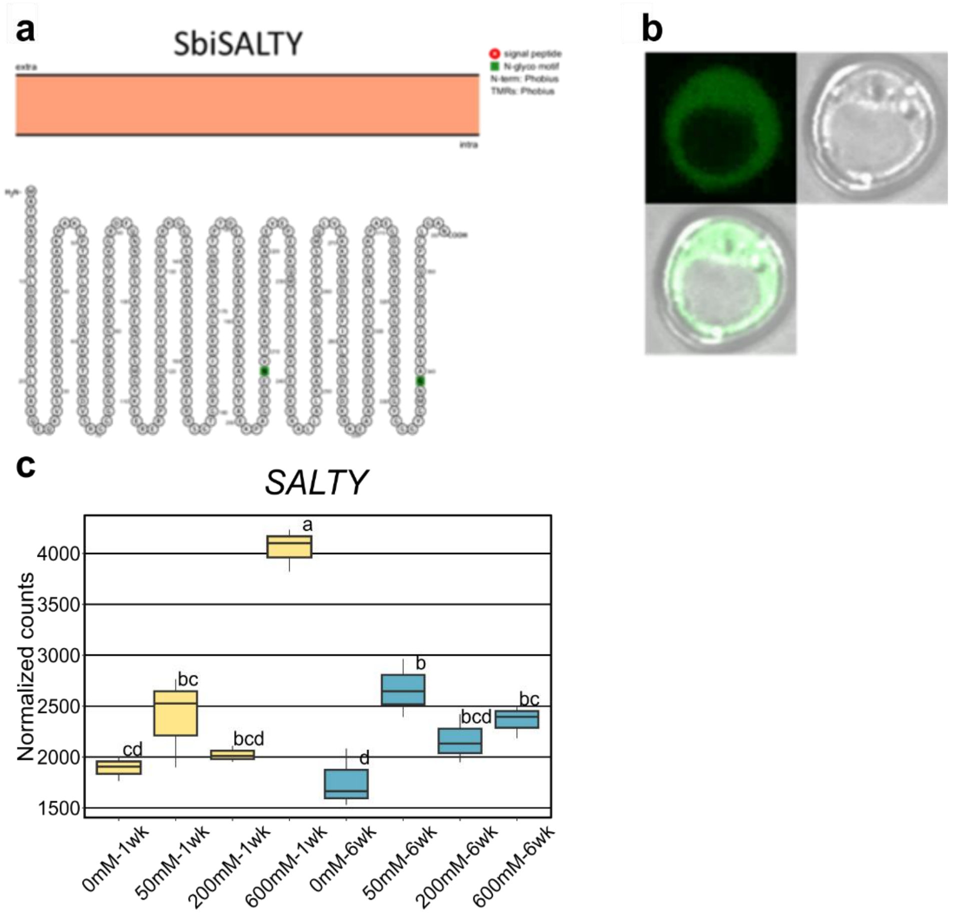


**Supplementary Figure 15.** Secondary structure and subcellular localization of SbiSALTY in yeast. **a,** Predicted secondary structure with Protter. No transmembrane helices are predicted for this protein. **b,** Subcellular localization observed by EGFP fusions in AXT3. The protein seems to localize to the cytosol. **c,** *SbiSALTY* gene expression in shoots of *S. bigelovii* plants treated with 0, 50, 200, and 600 mM NaCl for 1 and 6 weeks. Yellow, plants treated for 1 week; Blue, plants treated for 6 weeks.

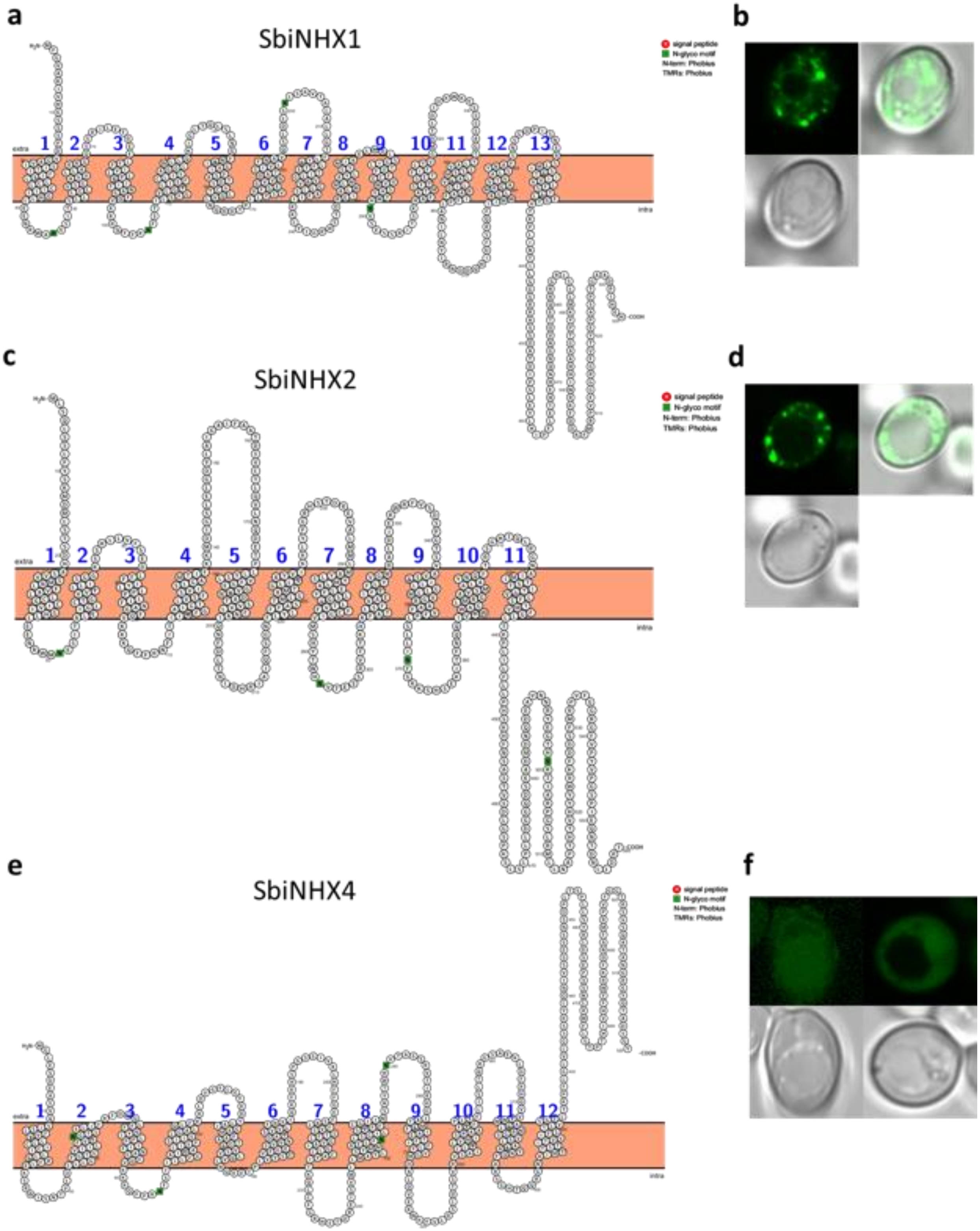
a
SbiNHX1
b
c
SbiNHX2
d
e
SbiNHX4
f
signal peptide
N-glyco motif
N-term: Phobius
TMRs: Phobius
extra
intra
-COOH

**Supplementary Figure 16.** Secondary structure and subcellular localization of SbiNHX proteins in yeast. Predicted secondary structure with Protter of: **a,** SbiNHX1; **c,** SbiNHX2; and **e,** SbiNHX4. Subcellular localization observed by EGFP fusions in AXT3 of: **b,** SbiNHX1; **d,** SbiNHX2; and **f**, SbiNHX4. SbiNHX1 and SbiNHX2 seem to be localized to lysosomes, while SbiNHX4 seems to have a vacuolar and ER localization.

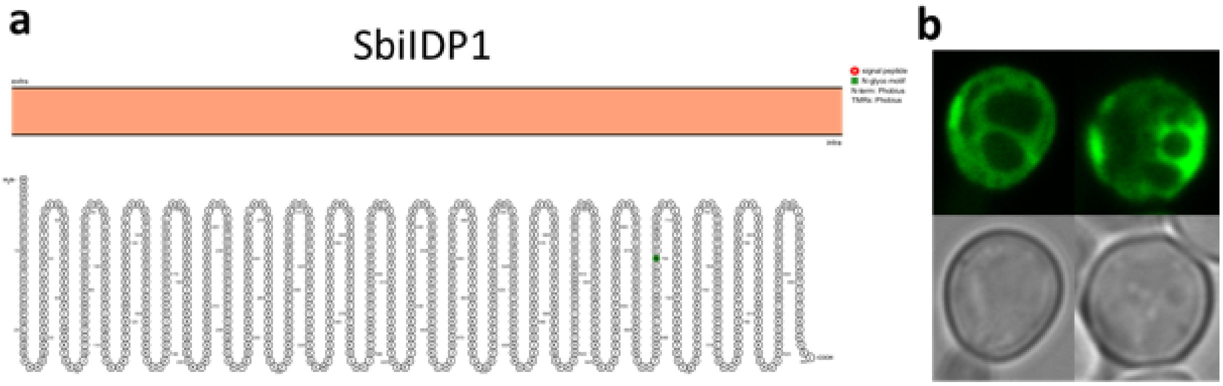


**Supplementary Figure 17.** Secondary structure and subcellular localization of SbiIDP1 in yeast. **a,** Predicted secondary structure with Protter; **b,** Subcellular localization observed by EGFP fusions in AXT3.

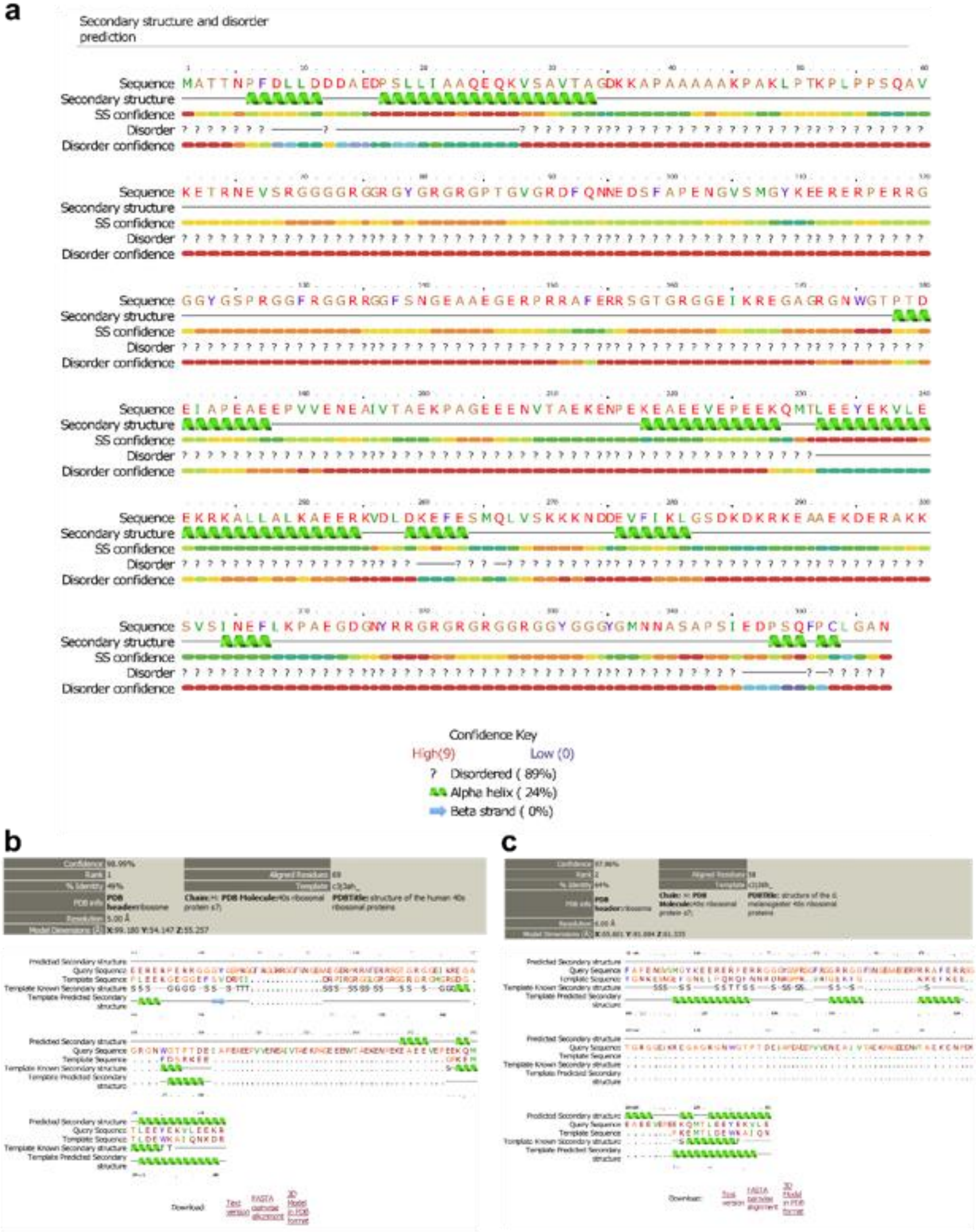
a
Secondary structure and disorder prediction
Sequence
Secondary structure
SS confidence
Disorder
Disorder confidence
Confidence Key
High(9)
Low (0)
? Disordered ( 89%)
Alpha helix ( 24%)
Beta strand ( 0%)
b
c
Predicted Secondary structure
Query Sequence
Template Sequence
Template Known Secondary structure
Template Predicted Secondary structure
Download
Text version
FASTA pairwise alignment
3D Model in PDB format

**Supplementary Figure 18.** SbiSALTY structure analysis with Phyre2. **a,** Secondary structure analysis of SbiSALTY. **b-c,** Best hits for homologous regions of SbiSALTY protein. **b,** Best hit against Homo sapiens 40S ribosomal protein S7. **c,** Second best hit against Drosophila melanogaster 40S ribosomal protein S7.
