## Supplementary Tables for "Learning from the expert: studying *Salicornia* to understand salinity tolerance"

**Supplementary Table 1.** ***Salicornia*** **genomes assembly statistics and completeness**

**Genome statistics**

| Species | Assembly size (Mb) | Scaffold number | Scaffold N50 (Mb) | Gap content (Ns %) | GC content (%) |
|---|---|---|---|---|---|
| *S. bigelovii* | 2,026.6 | 97 | 98.0 | 0.0 | 36.9 |
| *S. europaea* | 516.2 | 36 | 51.9 | 0.0 | 34.8 |

Annotation

| Species | Gene number | Isoform number | Repetitive elements (%) | Mean gene length (bp) | Mean cds length (bp) |
|---|---|---|---|---|---|
| *S. bigelovii* | 63,843 | 67,347 | 81.1 | 4,812 | 976 |
| *S. europaea* | 28,794 | 30,359 | 66.8 | 5,104 | 1,060 |

Annotation completeness (%) of Amaranthaceae genomes by BUSCO against the Eudicotyledons v10 dataset of 2,326 proteins, based on one isoform per gene

| Species | Complete | Complete single | Complete duplicated | Fragmented | Missing |
|---|---|---|---|---|---|
| *S. bigelovii* | 96.3 | 24.6 | 71.7 | 0.6 | 3.1 |
| *S. europaea* | 95.3 | 87.3 | 8.0 | 1.1 | 3.6 |
| *Beta vulgaris* | 96.2 | 85.9 | 10.3 | 1.0 | 2.8 |
| *Chenopodium quinoa* | 92.8 | 26.1 | 66.7 | 2.8 | 4.4 |
| *Spinacia oleracea* | 92.8 | 89.3 | 3.5 | 1.9 | 5.3 |

**Supplementary Table 2.** Matrix of the number of differentially expressed genes between treatments.

| | 1wk 0mM | 1wk 50mM | 1wk 200mM | 1wk 600mM | 6wk 0mM | 6wk 50mM | 6wk 200mM | 6wk 600mM |
|---|---|---|---|---|---|---|---|---|
| 1wk 0mM | - | | | | | | | |
| 1wk 50mM | 789 | - | | | | | | |
| 1wk 200mM | 618 | 80 | - | | | | | |
| 1wk 600mM | 5,486 | 2,857 | 2,838 | - | | | | |
| 6wk 0mM | 288 | 2,378 | 1,911 | 6,403 | - | | | |
| 6wk 50mM | 1,144 | 79 | 287 | 3,082 | 1,764 | - | | |
| 6wk 200mM | 1,438 | 960 | 645 | 4,564 | 808 | 172 | - | |
| 6wk 600mM | 1,866 | 604 | 1,501 | 5,024 | 1,788 | 103 | 8 | - |

**Supplementary Table 3.** Overrepresented MapMan3 bins for differentially expressed genes from shoots of *S. bigelovii*. Overrepresentation was tested with Wilcoxon Rank Sum test and corrected for multiple testing with Benjamini Hochberg with *p*-value < 0.05.

| **Bin** | **Description** | **Elements** | ***p*-value** |
|---|---|---|---|
| Treatment: 0 mM 1wk vs 50 mM 1wk NaCl | | | |
| 34.13 | transport peptides and oligopeptides | 8 | 0.02 |
| 10.6 | cell wall degradation | 7 | 0.02 |
| 26.10 | misc. cytochrome P450 | 6 | 0.03 |
| 10.5.1 | cell wall cell wall proteins AGPs | 7 | 0.03 |
| 10.5.1.1 | cell wall cell wall proteins AGPs AGP | 7 | 0.03 |
| 27.3.6 | RNA regulation of transcription bHLH Basic Helix-Loop-Helix family | 7 | 0.03 |
| Treatment: 0 mM 1wk vs 200 mM 1wk NaCl | | | |
| 10 | cell wall | 44 | 6.9E-4 |
| 16 | secondary metabolism | 28 | 1.0E-3 |
| 10.7 | cell wall modification | 6 | 7.0E-3 |
| 17.7 | hormone metabolism jasmonate | 11 | 7.0E-3 |
| 16.1 | secondary metabolism isoprenoids | 12 | 7.3E-3 |
| 10.5 | cell wall cell wall proteins | 11 | 0.01 |
| 17.7.1 | hormone metabolism jasmonate synthesis-degradation | 8 | 0.02 |
| 16.8 | secondary metabolism flavonoids | 5 | 0.04 |
| Treatment: 0 mM 1wk vs 600 mM 1wk NaCl | | | |
| 10 | cell wall | 122 | 1.0E-26 |
| 30.2.17 | signalling receptor kinases DUF 26 | 38 | 2.1E-12 |
| 29.2 | protein synthesis | 148 | 2.3E-10 |
| 29.2.1 | protein synthesis ribosomal protein | 74 | 1.9E-9 |
| 29 | protein | 744 | 2.4E-8 |
| 29.2.1.2 | protein synthesis ribosomal protein eukaryotic | 59 | 4.8E-8 |

| | | | |
|---|---|---|---|
| 29.5.11.4.4 | protein degradation ubiquitin E3 APC | 16 | 1.7E-7 |
| 10.5 | cell wall cell wall proteins | 19 | 2.9E-7 |
| 10.7 | cell wall modification | 16 | 6.9E-6 |
| 31 | cell | 181 | 8.4E-6 |
| 10.6 | cell wall degradation | 25 | 1.8E-5 |
| 11 | lipid metabolism | 109 | 1.8E-5 |
| 29.3 | protein targeting | 89 | 2.1E-5 |
| 26 | misc | 254 | 2.3E-5 |
| 26.12 | misc peroxidases | 18 | 2.3E-5 |
| 26.21 | misc protease inhibitor/seed storage/lipid transfer protein (LTP) family protein | 10 | 2.6E-5 |
| 27 | RNA | 495 | 2.6E-5 |
| 30.2.3 | signalling receptor kinases leucine rich repeat III | 16 | 6.3E-5 |
| 10.5.1 | cell wall cell wall proteins AGPs | 9 | 1.1E-4 |
| 10.5.1.1 | cell wall cell wall proteins AGPs AGP | 9 | 1.1E-4 |
| 10.2 | cell wall cellulose synthesis | 16 | 1.8E-4 |
| 29.2.1.2.2 | protein synthesis ribosomal protein eukaryotic 60S subunit | 32 | 2.1E-4 |
| 10.1 | cell wall precursor synthesis | 23 | 4.2E-4 |
| 9 | mitochondrial electron transport / ATP synthesis | 35 | 4.3E-4 |
| 11.1 | lipid metabolism FA synthesis and FA elongation | 32 | 4.5E-4 |
| 34 | transport | 229 | 6.0E-4 |
| 29.2.1.2.1 | protein synthesis ribosomal protein eukaryotic 40S subunit | 27 | 6.6E-4 |
| 34.19 | transport Major Intrinsic Proteins | 10 | 2.0E-3 |
| 26.2 | misc UDP glucosyl and glucoronyl transferases | 48 | 2.3E-3 |
| 27.3 | RNA regulation of transcription | 327 | 2.4E-3 |
| 30.5 | signalling G-proteins | 47 | 2.4E-3 |
| 31.4 | cell vesicle transport | 44 | 2.4E-3 |
| 31.1 | cell organisation | 81 | 2.5E-3 |
| 29.3.4 | protein targeting secretory pathway | 40 | 0.01 |
| 31.1.1 | cell organisation cytoskeleton | 26 | 0.01 |

| | | | |
|---|---|---|---|
| 10.6.3 | cell wall degradation pectate lyases and polygalacturonases | 13 | 0.02 |
| 27.4 | RNA RNA binding | 67 | 0.02 |
| 15.2 | metal handling binding, chelation and storage | 6 | 0.02 |
| 27.3.8 | RNA regulation of transcription C2C2(Zn) DOF zinc finger family | 5 | 0.02 |
| 30.2.11 | signalling receptor kinases leucine rich repeat XI | 7 | 0.02 |
| 31.1.1.2 | cell organisation cytoskeleton mikrotubuli | 14 | 0.02 |
| 20.2 | stress abiotic | 94 | 0.02 |
| 10.6.2 | cell wall degradation mannan-xylose-arabinose-fucose | 8 | 0.02 |
| 34.19.2 | transport Major Intrinsic Proteins TIP | 4 | 0.02 |
| 26.19 | misc plastocyanin-like | 6 | 0.02 |
| 30.2.24 | signalling receptor kinases S-locus glycoprotein like | 14 | 0.03 |
| 31.1.1.1.1 | cell organisation cytoskeleton actin Actin | 4 | 0.03 |
| 35.1.41 | not assigned no ontology hydroxyproline rich proteins | 13 | 0.03 |
| 11.9.3.2 | lipid metabolism lipid degradation lysophospholipases carboxylesterase | 5 | 0.03 |
| 11.9.3 | lipid metabolism lipid degradation lysophospholipases | 15 | 0.04 |
| 16.10 | secondary metabolism simple phenols | 5 | 0.04 |
| 34.8 | transport metabolite transporters at the envelope membrane | 15 | 0.04 |
| 35.1.19 | not assigned no ontology C2 domain-containing protein | 9 | 0.04 |
| 10.1.6 | cell wall precursor synthesis GAE | 5 | 0.04 |
| 35.1.5 | not assigned no ontology pentatricopeptide (PPR) repeat-containing protein | 85 | 0.04 |
| 21.4 | redox glutaredoxins | 5 | 0.04 |
| 13.1.3.4 | amino acid metabolism synthesis aspartate family methionine | 6 | 0.04 |
| 20.2.99 | stress abiotic unspecified | 12 | 0.04 |

| | | | |
|---|---|---|---|
| 34.3 | transport amino acids | 17 | 0.04 |
| 17.8 | hormone metabolism salicylic acid | 8 | 0.04 |
| 17.8.1 | hormone metabolism salicylic acid synthesis-degradation | 8 | 0.04 |
| 10.2.1 | cell wall cellulose synthesis cellulose synthase | 7 | 0.04 |
| 17.7.1.2 | hormone metabolism jasmonate synthesis-degradation lipoxygenase | 6 | 0.04 |
| 29.4.1 | protein postranslational modification kinase | 54 | 0.04 |
| 10.5.3 | cell wall cell wall proteins LRR | 4 | 0.04 |
| 33 | development | 134 | 0.04 |
| Treatment: 50 mM 1wk vs 200 mM 1wk NaCl | | | |
| 29.2 | protein synthesis | 46 | 5.3E-3 |
| 29.2.1 | protein synthesis ribosomal protein | 43 | 0.01 |
| 29.2.1.2 | protein synthesis ribosomal protein eukaryotic | 43 | 0.01 |
| 29.2.1.2.2 | protein synthesis ribosomal protein eukaryotic 60S subunit | 24 | 0.01 |
| Treatment: 50 mM 1wk vs 600 mM 1wk NaCl | | | |
| 10 | cell wall | 60 | 2.3E-9 |
| 27 | RNA | 269 | 5.1E-8 |
| 30.2.17 | signalling receptor kinases DUF 26 | 23 | 6.3E-8 |
| 26 | misc | 97 | 7.0E-6 |
| 9 | mitochondrial electron transport / ATP synthesis | 27 | 2.2E-5 |
| 34 | transport | 110 | 7.9E-5 |
| 27.3 | RNA regulation of transcription | 170 | 1.0E-4 |
| 29.5.11.4.4 | protein degradation ubiquitin E3 APC | 11 | 1.0E-4 |
| 29 | protein | 407 | 1.4E-4 |
| 26.2 | misc UDP glucosyl and glucoronyl transferases | 20 | 4.5E-4 |
| 9.1.2 | mitochondrial electron transport / ATP synthesis NADH-DH localisation not clear | 9 | 5.5E-4 |
| 9.1 | mitochondrial electron transport / ATP synthesis NADH-DH | 10 | 1.3E-3 |
| 33 | development | 58 | 1.8E-3 |

| | | | |
|---|---|---|---|
| 31 | cell | 84 | 1.8E-3 |
| 10.5 | cell wall cell wall proteins | 11 | 2.9E-3 |
| 29.5 | protein degradation | 170 | 2.9E-3 |
| 11 | lipid metabolism | 51 | 4.7E-3 |
| 30.2.3 | signalling receptor kinases leucine rich repeat III | 10 | 7.9E-3 |
| 29.3 | protein targeting | 55 | 9.4E-3 |
| 10.2 | cell wall cellulose synthesis | 10 | 0.01 |
| 33.99 | development unspecified | 52 | 0.01 |
| 31.4 | cell vesicle transport | 28 | 0.02 |
| 10.1 | cell wall precursor synthesis | 12 | 0.03 |
| 31.1.1 | cell organisation cytoskeleton | 15 | 0.04 |
| Treatment: 200 mM 1wk vs 600 mM 1wk NaCl | | | |
| 29.2 | protein synthesis | 124 | 4.0E-24 |
| 29 | protein | 437 | 4.4E-22 |
| 29.2.1 | protein synthesis ribosomal protein | 70 | 1.0E-17 |
| 29.2.1.2 | protein synthesis ribosomal protein eukaryotic | 59 | 2.1E-17 |
| 30.2.17 | signalling receptor kinases DUF 26 | 35 | 8.7E-14 |
| 27 | RNA | 230 | 3.2E-11 |
| 29.2.1.2.1 | protein synthesis ribosomal protein eukaryotic 40S subunit | 26 | 5.7E-9 |
| 10 | cell wall | 56 | 1.3E-8 |
| 29.2.1.2.2 | protein synthesis ribosomal protein eukaryotic 60S subunit | 33 | 3.0E-8 |
| 31 | cell | 88 | 2.3E-6 |
| 29.3 | protein targeting | 56 | 2.4E-6 |
| 26 | misc | 72 | 3.3E-6 |
| 27.4 | RNA RNA binding | 42 | 1.1E-5 |
| 20 | stress | 70 | 2.4E-5 |
| 34 | transport | 106 | 2.4E-5 |
| 30.2 | signalling receptor kinases | 80 | 5.3E-5 |
| 27.3 | RNA regulation of transcription | 132 | 5.9E-5 |
| 11 | lipid metabolism | 43 | 1.3E-4 |

| | | | |
|---|---|---|---|
| 29.5.11.4.4 | protein degradation ubiquitin E3 APC | 13 | 1.9E-4 |
| 29.6 | protein folding | 14 | 2.5E-4 |
| 20.2 | stress abiotic | 47 | 2.7E-4 |
| 13 | amino acid metabolism | 28 | 1.1E-3 |
| 13.1 | amino acid metabolism synthesis | 25 | 1.3E-3 |
| 26.2 | misc UDP glucosyl and glucoronyl transferases | 15 | 1.3E-37 |
| 10.1 | cell wall precursor synthesis | 14 | 1.8E-3 |
| 30.5 | signalling G-proteins | 25 | 2.4E-3 |
| 31.1.1 | cell organisation cytoskeleton | 14 | 4.6E-3 |
| 31.1 | cell organisation | 36 | 6.2E-3 |
| 29.3.4 | protein targeting secretory pathway | 22 | 6.3E-3 |
| 17.3 | hormone metabolism brassinosteroid | 7 | 6.4E-3 |
| 29.2.3 | protein synthesis initiation | 24 | 9.6E-3 |
| 8 | TCA / org transformation | 19 | 9.6E-3 |
| 29.2.4 | protein synthesis elongation | 10 | 0.01 |
| 10.7 | cell wall modification | 6 | 0.01 |
| 20.2.1 | stress abiotic heat | 20 | 0.01 |
| 34.8 | transport metabolite transporters at the envelope membrane | 11 | 0.01 |
| 29.2.2.3 | protein synthesis ribosome biogenesis Pre-rRNA processing and modifications | 13 | 0.01 |
| 31.4 | cell vesicle transport | 23 | 0.01 |
| 34.9 | transport metabolite transporters at the mitochondrial membrane | 14 | 0.01 |
| 29.4.1 | protein postranslational modification kinase | 29 | 0.01 |
| 11.1 | lipid metabolism FA synthesis and FA elongation | 19 | 0.01 |
| 27.1 | RNA processing | 56 | 0.01 |
| 7 | OPP | 7 | 0.01 |
| 29.5.11.20 | protein degradation ubiquitin proteasom | 17 | 0.01 |
| 21 | redox | 18 | 0.01 |
| 35.1.40 | not assigned no ontology glycine rich proteins | 9 | 0.01 |
| 11.9.3 | lipid metabolism lipid degradation lysophospholipases | 9 | 0.01 |

| | | | |
|---|---|---|---|
| 29.3.1 | protein targeting nucleus | 16 | 0.01 |
| 4 | glycolysis | 18 | 0.02 |
| 10.6 | cell wall degradation | 10 | 0.02 |
| 33 | development | 62 | 0.02 |
| 17.3.1 | hormone metabolism brassinosteroid synthesis-degradation | 4 | 0.02 |
| 31.1.1.2 | cell organisation cytoskeleton mikrotubuli | 9 | 0.02 |
| 4.1 | glycolysis cytosolic branch | 13 | 0.02 |
| 9 | mitochondrial electron transport / ATP synthesis | 14 | 0.03 |
| 30.2.3 | signalling receptor kinases leucine rich repeat III | 8 | 0.03 |
| 29.2.2 | protein synthesis ribosome biogenesis | 19 | 0.04 |
| 31.1.1.1.1 | cell organisation cytoskeleton actin Actin | 3 | 0.04 |
| 20.2.99 | stress abiotic unspecified | 5 | 0.04 |
| 27.3.99 | RNA regulation of transcription unclassified | 32 | 0.04 |
| Treatment: 0 mM 6wk vs 50 mM 6wk NaCl | | | |
| 28 | DNA | 46 | 6.3E-4 |
| 28.1 | DNA synthesis/chromatin structure | 34 | 1.5E-3 |
| 10 | cell wall | 62 | 5.4E-3 |
| 28.1.3 | DNA synthesis/chromatin structure histone | 13 | 5.4E-3 |
| 28.1.3.2 | DNA synthesis/chromatin structure histone core | 13 | 5.4E-3 |
| 30.2.17 | signalling receptor kinases DUF 26 | 9 | 0.02 |
| Treatment: 0 mM 6wk vs 200 mM 6wk NaCl | | | |
| 10 | cell wall | 56 | 2.2E-4 |
| 34.12 | transport metal | 10 | 0.04 |
| 10.6 | cell wall degradation | 13 | 0.04 |
| Treatment: 0 mM 6wk vs 600 mM 6wk NaCl | | | |
| 10 | cell wall | 60 | 4.2E-5 |
| 30.2.25 | signalling receptor kinases wall associated kinase | 6 | 9.8E-3 |
| 29.5 | protein degradation | 87 | 9.8E-3 |
| 10.5 | cell wall cell wall proteins | 8 | 9.8E-3 |

| | | | |
|---|---|---|---|
| 27.3.32 | RNA regulation of transcription WRKY domain transcription factor family | 8 | 0.02 |
| 1 | PS | 36 | 0.03 |
| 28.1.3.2 | DNA synthesis/chromatin structure histone core | 12 | 0.04 |
| Treatment: 50 mM 6wk vs 200 mM 6wk NaCl | | | |
| 20 | stress | 19 | 0.04 |
| Treatment: 50 mM 6wk vs 600 mM 6wk NaCl | | | |
| *NS* | | | |
| Treatment: 200 mM 6wk vs 600 mM 6wk NaCl | | | |
| *NS* | | | |

**Supplementary Table 4.** Overrepresented MapMan4 bins for differentially expressed genes from shoots of *S. bigelovii*. Overrepresentation was tested with Wilcoxon Rank Sum test and corrected for multiple testing with Benjamini Hochberg with *p*-value < 0.05.

| Bin | Description | Elements | *p*-value |
|---|---|---|---|
| Treatment: 0 mM 1wk vs 50 mM 1wk NaCl | | | |
| 25.4 | Nutrient uptake transition metal homeostasis | 11 | 3.7E-3 |
| 21.4 | Cell wall organisation cell wall proteins | 14 | 4.2E-3 |
| 21 | Cell wall organisation | 42 | 4.2E-3 |
| 25 | Nutrient uptake | 14 | 4.2E-3 |
| 9 | Secondary metabolism | 13 | 4.2E-3 |
| 21.4.1 | Cell wall organisation cell wall proteins hydroxyproline-rich glycoprotein activities | 12 | 6.5E-3 |
| 21.4.1.1 | Cell wall organisation cell wall proteins hydroxyproline-rich glycoprotein activities arabinogalactan-protein activities | 9 | 7.3E-3 |

| | | | |
|---|---|---|---|
| 21.4.1.1.3 | Cell wall organisation cell wall proteins hydroxyproline-rich glycoprotein activities arabinogalactan-protein activities Fasciclin-type arabinogalactan protein (FLA) | 7 | 0.04 |
| Treatment: 0 mM 1wk vs 200 mM 1wk NaCl | | | |
| 9 | Secondary metabolism | 16 | 3.8E-3 |
| 21.4 | Cell wall organisation cell wall proteins | 12 | 5.7E-3 |
| 21 | Cell wall organisation | 48 | 5.7E-3 |
| 9.1 | Secondary metabolism terpenoids | 13 | 0.01 |
| 21.4.1 | Cell wall organisation cell wall proteins hydroxyproline-rich glycoprotein activities | 11 | 0.01 |
| Treatment: 0 mM 1wk vs 600 mM 1wk NaCl | | | |
| 21 | Cell wall organisation | 149 | 3.9E-27 |
| 17 | Protein biosynthesis | 184 | 5.7E-13 |
| 21.4 | Cell wall organisation cell wall proteins | 27 | 6.1E-12 |
| 17.1 | Protein biosynthesis ribosome biogenesis | 89 | 2.9E-10 |
| 21.4.1 | Cell wall organisation cell wall proteins hydroxyproline-rich glycoprotein activities | 21 | 1.7E-9 |
| 21.4.1.1 | Cell wall organisation cell wall proteins hydroxyproline-rich glycoprotein activities arabinogalactan-protein activities | 18 | 2.9E-8 |
| 5 | Lipid metabolism | 117 | 2.0E-6 |
| 18 | Protein modification | 289 | 4.0E-6 |
| 21.1 | Cell wall organisation cellulose | 25 | 1.3E-5 |
| 22 | Vesicle trafficking | 125 | 1.3E-5 |
| 18.4.1 | Protein modification phosphorylation TKL protein kinase superfamily | 121 | 2.1E-5 |
| 21.2 | Cell wall organisation hemicellulose | 29 | 5.5E-5 |
| 18.4 | Protein modification phosphorylation | 221 | 7.8E-5 |
| 21.4.1.1.3 | Cell wall organisation cell wall proteins hydroxyproline-rich glycoprotein activities arabinogalactan-protein activities Fasciclin-type arabinogalactan protein (FLA) | 10 | 1.5E-4 |

| | | | |
|---|---|---|---|
| 17.1.3 | Protein biosynthesis ribosome biogenesis small ribosomal subunit (SSU) | 36 | 2.1E-4 |
| 24.2 | Solute transport carrier-mediated transport | 149 | 2.4E-4 |
| 5.1 | Lipid metabolism fatty acid metabolism | 48 | 3.3E-4 |
| 23 | Protein translocation | 63 | 5.4E-4 |
| 21.1.1 | Cell wall organisation cellulose cellulose synthase complex (CSC) | 17 | 6.9E-4 |
| 11.10 | Phytohormone action signalling peptides | 29 | 9.1E-4 |
| 11 | Phytohormone action | 102 | 9.1E-4 |
| 17.1.2 | Protein biosynthesis ribosome biogenesis large ribosomal subunit (LSU) | 37 | 9.1E-4 |
| 11.10.2 | Phytohormone action signalling peptides CRP (cysteine-rich-peptide) category | 20 | 1.0E-3 |
| 17.1.2.1 | Protein biosynthesis ribosome biogenesis large ribosomal subunit (LSU) LSU proteome | 26 | 1.3E-3 |
| 3.13 | Carbohydrate metabolism nucleotide sugar biosynthesis | 26 | 1.4E-3 |
| 17.1.3.1 | Protein biosynthesis ribosome biogenesis small ribosomal subunit (SSU) SSU proteome | 27 | 1.5E-3 |
| 2 | Cellular respiration | 54 | 1.5E-3 |
| 24 | Solute transport | 241 | 2.6E-3 |
| 5.5 | Lipid metabolism phytosterol metabolism | 11 | 3.4E-3 |
| 50.2.4 | Enzyme classification EC_2 transferases EC_2.4 glycosyltransferase | 23 | 3.8E-3 |
| 3 | Carbohydrate metabolism | 83 | 3.8E-3 |
| 2.4 | Cellular respiration oxidative phosphorylation | 32 | 5.0E-3 |
| 20.1.1 | Cytoskeleton organisation microtubular network alpha-beta-Tubulin heterodimer | 8 | 5.5E-3 |
| 13.2.4 | Cell division cell cycle organisation metaphase to anaphase transition | 9 | 5.8E-3 |
| 13.2.4.1 | Cell division cell cycle organisation metaphase to anaphase transition Anaphase-Promoting Complex/Cyclosome (APC/C)-dependent ubiquitination | 9 | 5.8E-3 |

| | | | |
|---|---|---|---|
| 13.2.4.1.1 | Cell division cell cycle organisation metaphase to anaphase transition Anaphase-Promoting Complex/Cyclosome (APC/C)-dependent ubiquitination APC/C E3 ubiquitin protein ligase complex | 9 | 5.8E-3 |
| 20 | Cytoskeleton organisation | 80 | 6.5E-3 |
| 13.2.4.1.1.1 | Cell division cell cycle organisation metaphase to anaphase transition Anaphase-Promoting Complex/Cyclosome (APC/C)-dependent ubiquitination APC/C E3 ubiquitin protein ligase complex platform subcomplex | 7 | 7.2E-3 |
| 13.2.4.1.1.1.1 | Cell division cell cycle organisation metaphase to anaphase transition Anaphase-Promoting Complex/Cyclosome (APC/C)-dependent ubiquitination APC/C E3 ubiquitin protein ligase complex platform subcomplex component APC1 | 7 | 7.2E-3 |
| 23.5 | Protein translocation nucleus | 30 | 0.01 |
| 18.4.1.3 | Protein modification phosphorylation TKL protein kinase superfamily protein kinase (LRR-III) | 14 | 0.01 |
| 13.1 | Cell division DNA replication | 31 | 0.01 |
| 21.3 | Cell wall organisation pectin | 41 | 0.01 |
| 21.6.2 | Cell wall organisation lignin monolignol conjugation and polymerization | 6 | 0.01 |
| 24.3.1 | Solute transport channels MIP family | 9 | 0.01 |
| 20.1 | Cytoskeleton organisation microtubular network | 27 | 0.01 |
| 11.10.2.1 | Phytohormone action signalling peptides CRP (cysteine-rich-peptide) category GASA/GAST-peptide activity | 11 | 0.02 |
| 11.10.2.1.1 | Phytohormone action signalling peptides CRP (cysteine-rich-peptide) category GASA/GAST-peptide activity GASA-precursor polypeptide | 11 | 0.02 |
| 16 | RNA processing | 190 | 0.02 |
| 21.2.2 | Cell wall organisation hemicellulose xylan | 9 | 0.02 |

| | | | |
|---|---|---|---|
| 23.5.2 | Protein translocation nucleus nucleocytoplasmic transport | 15 | 0.02 |
| 5.5.1 | Lipid metabolism phytosterol metabolism plant sterol pathway | 6 | 0.02 |
| 50.1.13 | Enzyme classification EC_1 oxidoreductases EC_1.14 oxidoreductase acting on paired donor with incorporation or reduction of molecular oxygen | 31 | 0.02 |
| 15.5 | RNA biosynthesis transcriptional regulation | 216 | 0.03 |
| 22.2 | Vesicle trafficking retrograde trafficking | 25 | 0.03 |
| 21.4.2 | Cell wall organisation cell wall proteins expansin activities | 6 | 0.03 |
| 16.4.9 | RNA processing RNA homeostasis mRNA stress granule formation | 18 | 0.03 |
| 22.5.2.4.3 | Vesicle trafficking multi-pathway trafficking regulation vesicle tethering RAB-GTPase membrane association RAB-GDI displacement factor (GDF) activities | 7 | 0.03 |
| 22.5.2.4.3.2 | Vesicle trafficking multi-pathway trafficking regulation vesicle tethering RAB-GTPase membrane association RAB-GDI displacement factor (GDF) activities B-G-class Rab-GDF protein | 6 | 0.03 |
| 22.5 | Vesicle trafficking multi-pathway trafficking regulation | 69 | 0.04 |
| 21.2.2.1 | Cell wall organisation hemicellulose xylan biosynthesis | 7 | 0.04 |
| 19.2.2.1.4.3.3 | Protein homeostasis ubiquitin-proteasome system ubiquitin-fold protein conjugation ubiquitin conjugation (ubiquitylation) ubiquitin-ligase E3 activities RING-domain E3 ligase activities RING-H2-class ligase activities | 16 | 0.04 |
| 19.2.2.1.4.3.3.1 | Protein homeostasis ubiquitin-proteasome system ubiquitin-fold protein conjugation | 9 | 0.04 |

| | | | |
|---|---|---|---|
| | ubiquitin conjugation (ubiquitylation) ubiquitin-ligase E3 activities RING-domain E3 ligase activities RING-H2-class ligase activities ATL-subclass ligase | | |
| 24.2.4.1.1 | Solute transport carrier-mediated transport MOP superfamily MATE family metabolite transporter (DTX) | 5 | 0.04 |
| 15.5.1.5 | RNA biosynthesis transcriptional regulation C2C2 transcription factor superfamily transcription factor (DOF) | 5 | 0.04 |
| Treatment: 50 mM 1wk vs 200 mM 1wk NaCl | | | |
| 17 | Protein biosynthesis | 47 | 4.6E-3 |
| 17.1.2 | Protein biosynthesis ribosome biogenesis large ribosomal subunit (LSU) | 25 | 4.6E-3 |
| 17.1.2.1 | Protein biosynthesis ribosome biogenesis large ribosomal subunit (LSU) LSU proteome | 25 | 4.6E-3 |
| 17.1 | Protein biosynthesis ribosome biogenesis | 45 | 4.6E-3 |
| Treatment: 50 mM 1wk vs 600 mM 1wk NaCl | | | |
| 21 | Cell wall organisation | 78 | 1.7E-13 |
| 22 | Vesicle trafficking | 86 | 7.7E-6 |
| 19 | Protein homeostasis | 144 | 3.9E-5 |
| 24 | Solute transport | 115 | 4.6E-4 |
| 16 | RNA processing | 129 | 4.6E-4 |
| 18.4 | Protein modification phosphorylation | 103 | 4.7E-4 |
| 3 | Carbohydrate metabolism | 43 | 6.1E-4 |
| 18 | Protein modification | 141 | 6.4E-4 |
| 15 | RNA biosynthesis | 109 | 6.4E-4 |
| 21.1 | Cell wall organisation cellulose | 17 | 9.1E-4 |
| 18.4.1 | Protein modification phosphorylation TKL protein kinase superfamily | 59 | 1.2E-3 |
| 21.4 | Cell wall organisation cell wall proteins | 12 | 1.5E-3 |
| 15.5 | RNA biosynthesis transcriptional regulation | 82 | 5.7E-3 |

| | | | |
|---|---|---|---|
| 3.13 | Carbohydrate metabolism nucleotide sugar biosynthesis | 15 | 7.5E-3 |
| 21.2 | Cell wall organisation hemicellulose | 17 | 7.6E-3 |
| 21.1.1 | Cell wall organisation cellulose cellulose synthase complex (CSC) | 14 | 0.01 |
| 11 | Phytohormone action | 48 | 0.01 |
| 16.4 | RNA processing RNA homeostasis | 33 | 0.01 |
| 19.2 | Protein homeostasis ubiquitin-proteasome system | 87 | 0.02 |
| 24.2 | Solute transport carrier-mediated transport | 77 | 0.02 |
| 27 | Multi-process regulation | 43 | 0.02 |
| 21.4.1 | Cell wall organisation cell wall proteins hydroxyproline-rich glycoprotein activities | 9 | 0.02 |
| 12 | Chromatin organisation | 49 | 0.02 |
| 13.2.4.1.1 | Cell division cell cycle organisation metaphase to anaphase transition Anaphase-Promoting Complex/Cyclosome (APC/C)-dependent ubiquitination APC/C E3 ubiquitin protein ligase complex | 6 | 0.03 |
| 13.2.4.1.1.1 | Cell division cell cycle organisation metaphase to anaphase transition Anaphase-Promoting Complex/Cyclosome (APC/C)-dependent ubiquitination APC/C E3 ubiquitin protein ligase complex platform subcomplex | 6 | 0.03 |
| 13.2.4.1.1.1.1 | Cell division cell cycle organisation metaphase to anaphase transition Anaphase-Promoting Complex/Cyclosome (APC/C)-dependent ubiquitination APC/C E3 ubiquitin protein ligase complex platform subcomplex component APC1 | 6 | 0.03 |
| 5 | Lipid metabolism | 58 | 0.03 |
| 19.2.2.1 | Protein homeostasis ubiquitin-proteasome system ubiquitin-fold protein conjugation ubiquitin conjugation (ubiquitylation) | 28 | 0.04 |

| | | | |
|---|---|---|---|
| 16.4.9 | RNA processing RNA homeostasis mRNA stress granule formation | 13 | 0.04 |
| Treatment: 200 mM 1wk vs 600 mM 1wk NaCl | | | |
| 17 | Protein biosynthesis | 145 | 5.6E-26 |
| 17.1 | Protein biosynthesis ribosome biogenesis | 78 | 6.8E-21 |
| 17.1.2.1 | Protein biosynthesis ribosome biogenesis large ribosomal subunit (LSU) LSU proteome | 30 | 6.4E-9 |
| 17.1.3 | Protein biosynthesis ribosome biogenesis small ribosomal subunit (SSU) | 30 | 7.3E-9 |
| 17.1.2 | Protein biosynthesis ribosome biogenesis large ribosomal subunit (LSU) | 37 | 1.1E-8 |
| 17.1.3.1 | Protein biosynthesis ribosome biogenesis small ribosomal subunit (SSU) SSU proteome | 26 | 1.9E-8 |
| 16 | RNA processing | 108 | 7.5E-7 |
| 21 | Cell wall organisation | 63 | 2.7E-6 |
| 5 | Lipid metabolism | 57 | 5.0E-6 |
| 19 | Protein homeostasis | 129 | 5.0E-6 |
| 24 | Solute transport | 107 | 1.8E-5 |
| 16.4 | RNA processing RNA homeostasis | 29 | 5.2E-5 |
| 24.2 | Solute transport carrier-mediated transport | 69 | 1.0E-4 |
| 35.1 | not assigned annotated | 530 | 1.2E-4 |
| 2 | Cellular respiration | 37 | 1.3E-4 |
| 23 | Protein translocation | 41 | 1.3E-4 |
| 5.1 | Lipid metabolism fatty acid metabolism | 30 | 1.3E-4 |
| 22 | Vesicle trafficking | 64 | 2.6E-4 |
| 3 | Carbohydrate metabolism | 43 | 3.2E-4 |
| 21.4 | Cell wall organisation cell wall proteins | 9 | 8.0E-4 |
| 20 | Cytoskeleton organisation | 48 | 1.5E-3 |
| 3.13 | Carbohydrate metabolism nucleotide sugar biosynthesis | 15 | 2.6E-3 |
| 27 | Multi-process regulation | 32 | 2.6E-3 |
| 4 | Amino acid metabolism | 29 | 3.0E-3 |
| 18 | Protein modification | 130 | 3.9E-3 |

| | | | |
|---|---|---|---|
| 16.4.9 | RNA processing RNA homeostasis mRNA stress granule formation | 16 | 5.5E-3 |
| 19.2 | Protein homeostasis ubiquitin-proteasome system | 80 | 5.6E-3 |
| 2.4 | Cellular respiration oxidative phosphorylation | 22 | 7.0E-3 |
| 19.1 | Protein homeostasis protein quality control | 17 | 0.01 |
| 23.5 | Protein translocation nucleus | 19 | 0.03 |
| 17.6.1 | Protein biosynthesis organelle machinery mitochondrial ribosome biogenesis | 15 | 0.03 |
| 27.5 | Multi-process regulation ROP-GTPase regulatory system | 5 | 0.04 |
| 17.3 | Protein biosynthesis translation initiation | 26 | 0.04 |
| Treatment: 0 mM 6wk vs 50 mM 6wk NaCl | | | |
| 21 | Cell wall organisation | 81 | 0.03 |
| 12.1.1 | Chromatin organisation chromatin structure DNA wrapping | 13 | 0.03 |
| 12 | Chromatin organisation | 34 | 0.03 |
| 12.1 | Chromatin organisation chromatin structure | 18 | 0.03 |
| 50.2.4 | Enzyme classification EC_2 transferases EC_2.4 glycosyltransferase | 13 | 0.04 |
| Treatment: 0 mM 6wk vs 200 mM 6wk NaCl | | | |
| *NS* | | 56 | 2.2E-4 |
| Treatment: 0 mM 6wk vs 600 mM 6wk NaCl | | | |
| 50.2.4 | Enzyme classification EC_2 transferases EC_2.4 glycosyltransferase | 14 | 0.04 |
| 18.4.1.25 | Protein modification phosphorylation TKL protein kinase superfamily protein kinase (WAK/WAKL) | 6 | 0.04 |
| Treatment: 50 mM 6wk vs 200 mM 6wk NaCl | | | |
| *NS* | | | |
| Treatment: 50 mM 6wk vs 600 mM 6wk NaCl | | | |
| *NS* | | | |

| |
|---|
| Treatment: 200 mM 6wk vs 600 mM 6wk NaCl |
| *NS* |

**Supplementary Table 5.** Overrepresented MapMan3 bins for differentially abundant proteins from shoots of *S. bigelovii* by treatment. Overrepresentation was tested with Wilcoxon Rank Sum test and corrected for multiple testing with Benjamini Hochberg with p value < 0.05.

| **Bin** | **Description** | **Elements** | ***p*-value** |
|---|---|---|---|
| Treatment: 0 mM and 50 mM up vs 200 mM and 600 mM down | | | |
| 30 | Signaling | 27 | 6.6E-4 |
| Treatment: 0 mM and 50 mM down vs 200 mM and 600 mM up | | | |
| 9 | Mitochondrial electron transport / ATP synthesis | 26 | 8.3E-5 |

**Supplementary Table 6.** Overrepresented MapMan4 bins for differentially abundant proteins from shoots of *S. bigelovii* by treatment. Overrepresentation was tested with Wilcoxon Rank Sum test and corrected for multiple testing with Benjamini Hochberg with p value < 0.05.

| **Bin** | **Description** | **Elements** | ***p*-value** |
|---|---|---|---|
| Treatment: 0 mM and 50 mM up vs 200 mM and 600 mM down | | | |
| 22.5 | Vesicle trafficking multi-pathway trafficking regulation | 15 | 0.03 |
| 23.1 | Protein translocation chloroplast | 15 | 0.04 |
| Treatment: 0 mM and 50 mM down vs 200 mM and 600 mM up | | | |
| 2 | Cellular respiration | 37 | 1.6E-6 |
| 2.4 | Cellular respiration oxidative phosphorylation | 30 | 1.6E-6 |

| | | | |
|---|---|---|---|
| 2.4.3 | Cellular respiration oxidative phosphorylation cytochrome c reductase complex | 8 | 8.1E-3 |
| 17.1 | Protein biosynthesis ribosome biogenesis | 6 | 0.04 |

**Supplementary Table 7.** Genes affecting AXT3 growth under NaCl.

| Gene alias | Effect in salt tolerance | Description |
|---|---|---|
| *Sbi_SALTY* | Increase (+ + +) | Hyaluronan / mRNA binding |
| *Sbi_KAT2* | Increase (+ + +) | Potassium channel KAT2 |
| *Sbi_NHX4* | Increase (+ +) | Sodium/hydrogen exchanger |
| *Sbi_OPT4* | Increase (+) | Oligopeptide transporter |
| *Sbi_IDP1* | Decrease (- - -) | Unknown |
| *Sbi_HKT1* | Decrease (- - -) | Sodium transporter HKT1 |
| *Sbi_NHX1* | Decrease (- - -) | Sodium/hydrogen exchanger |
| *Sbi_NHX2* | Decrease (- - -) | Sodium/hydrogen exchanger |
| *Sbi_ENT1* | Decrease (- -) | Equilibrative nucleotide transporter |
| *Sbi_SLC5-6* | Decrease (- -) | Sodium-coupled neutral amino acid transporter |
| *Sbi_PQ* | Decrease (- -) | PQ-loop, Probable vacuolar amino acid transporter |
